# A genome-wide loss-of-function screen identifies *Toxoplasma gondii* genes that determine fitness in interferon gamma-activated murine macrophages

**DOI:** 10.1101/867705

**Authors:** Yifan Wang, Lamba Omar Sangaré, Tatiana C. Paredes-Santos, Shruthi Krishnamurthy, Musa A. Hassan, Anna M. Furuta, Benedikt M. Markus, Sebastian Lourido, Jeroen P.J. Saeij

## Abstract

Macrophages play an essential role in the early immune response against *Toxoplasma* and are the cell type preferentially infected by the parasite *in vivo*. Interferon gamma (IFNγ) elicits a variety of anti-*Toxoplasma* activities in macrophages. Using a genome-wide CRISPR screen we identified ∼130 *Toxoplasma* genes that determine parasite fitness in naїve macrophages and ∼466 genes that determine fitness in IFNγ-stimulated murine macrophages, seven of which we investigated and confirmed. We show that one of these genes encodes dense granule protein GRA45, which contains a putative chaperone-like domain, and which we show is critical in preventing other GRA effectors from aggregating. Parasites lacking *GRA45* mislocalize GRA effectors upon secretion, are more susceptible to IFNγ-mediated growth inhibition, and have reduced virulence in mice. Our results provide a resource for the community to further explore the function of *Toxoplasma* genes that determine fitness in IFNγ-stimulated macrophages.

**IMPORTANCE:** The intracellular parasite *Toxoplasma gondii* can cause congenital infections and severe disease in immunocompromised patients. The cytokine IFNγ can block parasite replication by upregulating a variety of toxoplasmacidal mechanisms in many cells, including macrophages. *Toxoplasma* preferentially infects macrophages. Therefore, the parasite has evolved mechanisms to survive in these cells in the presence of IFNγ. Here, we generated pools of *Toxoplasma* mutants for every gene and determined which mutants were specifically depleted in IFNγ-stimulated macrophages, thus identifying parasite genes determining fitness in these cells. We show that one of these genes encodes for a dense granule protein (GRA45) that plays an important role in preventing other GRA effectors from aggregating. Parasites without GRA45 mislocalize GRA effectors upon secretion, have enhanced susceptibility to IFNγ-mediated growth inhibition, and are avirulent in mice. Thus, our screen provides a resource to the *Toxoplasma* community to determine the function of *Toxoplasma* genes that affect its fitness in IFNγ-stimulated macrophages.

## INTRODUCTION

*Toxoplasma gondii* is an obligate intracellular parasite that causes disease in immunocompromised individuals, such as HIV/AIDS patients, and when contracted congenitally (1). It causes lifelong infections by converting from rapidly dividing tachyzoite stages into encysted slow-growing bradyzoites, which mainly localize to the brain and muscle tissues. Once established in the host cell, *Toxoplasm*a resides in a unique replication niche called the parasitophorous vacuole (PV), which is separated from the host cytoplasm by the PV membrane (PVM) and does not fuse with the endolysosomal system (2, 3).

The cytokine interferon gamma (IFNγ) is essential for the control of *Toxoplasma* replication in host cells (4). Mice with macrophages that can no longer respond to IFNγ are extremely susceptible to *Toxoplasma* demonstrating the crucial role of this cell type in IFNγ-mediated control of *Toxoplasma* (5). IFNγ-induced upregulation of immunity-related GTPases (IRGs) and guanylate binding proteins (GBPs) (6–8) are key mechanisms for controlling *Toxoplasma* in mice (9–13). IRGs and GBPs can bind and subsequently vesiculate the PVM, eventually leading to the death of the parasite (14–17). In addition, IFNγ induces the production of nitric oxide (NO) and reactive oxygen species, which lead to parasite damage and growth restriction in murine macrophages (18). However, the potency of IFNγ-induced toxoplasmacidal activities in murine macrophages is parasite-strain dependent, with the clonal type 2 and 3 strains being more susceptible to IFNγ-induced growth restriction compared to type 1 strains (19).

*Toxoplasma* co-opts host cells by secreting ROP and GRA effector proteins, from organelles called rhoptries and dense granules, respectively, into the host cell or onto the PVM (reviewed in (20)). *ROP18* (encoding a secreted kinase) (21, 22) and *ROP5* (encoding a member of an expanded family of pseudokinases) (23, 24) are highly polymorphic and, together with ROP17 (25) and GRA7 (26, 27), account for differences in parasite virulence in mice by cooperatively blocking IRG (14, 28–30) and GBP (6, 31) loading on the PVM. Other parasite strain-specific effectors include ROP16 and GRA15, which can affect GBP loading on the PVM (32) and macrophage polarization into the classical (M1) or alternative (M2) activation phenotypes (33). The pseudokinase ROP54 also inhibits GBP loading on the PVM (34). Additional secreted effectors, such as GRA12, affect *Toxoplasma* survival in murine macrophages without affecting the loading of IRGs/GBPs to the PVM (35). Besides targeting IFNγ-induced host cell effectors, *Toxoplasma* can directly inhibit IFNγ signaling. For example, through a GRA (*Tg*IST) secreted beyond the PVM (36, 37), *Toxoplasma* can directly inhibit STAT1 transcriptional activity (38–40), an essential component of the IFNγ signaling pathway (41). The secretion of GRAs beyond the PVM into the host cell cytosol is dependent on the *Toxoplasma* proteins MYR1/2/3 (42, 43), which might form a PVM-localized translocon. A Golgi-resident aspartyl protease ASP5, which cleaves the *Toxoplasma* export element (TEXEL, RRLxx) motif in some GRAs, is required for exported GRAs to reach the host cytosol and also for the correct localization of other GRAs confined to the PV (44–46).

Many of the above-mentioned *Toxoplasma* virulence genes were identified using genetic crosses between strains that differ in virulence in mice. However, this approach fails to identify *Toxoplasma* genes that determine strain-independent survival in IFNγ-activated macrophages. Here we used a genome-wide loss-of-function screen in the type 1 (RH) strain, which is resistant to IRG/GBP-mediated killing, to identify ∼466 *Toxoplasma* genes that determine fitness in IFNγ-activated murine macrophages and ∼130 genes that determine fitness in naїve macrophages. Seven of the top hits, including GRA45, were confirmed by generating and testing single gene knockouts in functional parasite growth assays in IFNγ-activated murine macrophages. We show that GRA45 plays an important role in preventing aggregation of other GRA effectors in the secretory pathway. Δ*gra45* parasites have enhanced susceptibility to IFNγ-mediated growth inhibition in murine, rat, and human macrophages and significantly reduced virulence in mice. Our results provide a resource to the community to investigate the mechanism by which other *Toxoplasma* genes affect fitness in naїve or activated macrophages.

## RESULTS

### A loss-of-function screen identifies *Toxoplasma* genes that determine fitness in naïve and IFNγ-activated murine macrophages

We previously identified *Toxoplasma* genes that determine parasite fitness during infection of human foreskin fibroblasts (HFFs) using a genome-wide loss-of-function CRISPR/Cas9 screen (47). However, it is likely that this set of fitness-conferring genes varies by host cell type, host species, and in infection of IFNγ-activated cells. The challenge in *Toxoplasma* genome-wide loss-of-function screens is to maintain the complexity of the mutant pool during the selection. It was previously reported that macrophages require stimulation by IFNγ along with LPS or TNFα to effectively restrict *Toxoplasma* growth (48). Such stimulated macrophages are extremely effective in inhibiting *Toxoplasma* growth, even of virulent strains like the type 1 (RH) strain, which would likely pose a tight bottleneck during the selection of the mutant parasite pool. IFNγ stimulation by itself was sufficient for a minor but significant inhibition of *Toxoplasma* growth in C57BL/6J murine bone marrow-derived macrophages (BMDMs) (**Figure S1A**), unlike IFNγ + TNFα stimulation that drastically restricted parasite growth (**Figure S1B**). Thus, we reasoned that a loss-of-function screen in macrophages stimulated with IFNγ alone would reduce the random loss of mutants while at the same time allowing us to identify mutants that are susceptible to IFNγ-mediated restriction of parasite growth. To identify these mutants, we applied a genome-wide CRISPR loss-of-function screen in RH parasites constitutively expressing Cas9 (RH-Cas9) (47). We initially passed the mutant pool in HFFs to remove mutants with general fitness defects (e.g., invasion and replication) (**Figure 1A**). Parasite mutants coming out of the 4^th^ HFF passage were subsequently passed three times on either naïve or IFNγ-stimulated BMDMs (100 ng/mL IFNγ for 24 h) (referred to as screen 1 or S1) or were continued to be passed on HFFs (**Figure 1A and 1B**). sgRNAs were amplified from the input library, and from parasite DNA isolated after 4^th^ and 8^th^ passages on HFFs and three additional passages on naïve or IFNγ-activated BMDMs and enumerated using Illumina sequencing. To assess potential bottlenecks and associated random loss of mutants at different passages, we enumerated sgRNAs targeting a set of control genes that are only expressed in sexual stages and should, therefore, have no fitness effect (49). The sgRNAs targeting these control genes were significantly enriched (*p* < 0.0001, chi-square) at later passages, confirming that they do not affect parasite fitness. However, we noted that there was still a discernible loss of sgRNAs targeting control genes in IFNγ-activated BMDMs (S1 in **Figure 1C**) indicating that some stochastic loss of library complexity occurs at this IFNγ concentration. Because an IFNγ concentration of 1 ng/mL induced ∼20% inhibition of parasite growth (**Figure S1A**), we repeated the screen in BMDMs pre-stimulated with 1 ng/mL IFNγ for 24 h (referred to as screen 2 or S2) (**Figure 1B**). In addition, to better understand the changes in stimulated BMDMs that *Toxoplasma* needs to adapt to, we performed RNA-sequencing (RNA-seq) on C57BL/6J and A/J BMDMs stimulated with IFNγ and/or TNFα for 4 h or 24 h. These analyses showed that IFNγ stimulation for 4 and 24 h induced the expression of similar gene sets (**Figure S1C and Table S1**) while TNFα-stimulated macrophages had different gene sets and profiles. In an attempt to reduce the IFNγ-induced bottleneck in the screen and to identify parasite genes important for fitness in IFNγ-stimulated BMDMs and specific for that stimulation, we performed a 3^rd^ screen in BMDMs stimulated with IFNγ or TNFα for 4 h before infection with the pool of mutants coming out of the 3^rd^ passage on HFFs (referred to as screen 3 or S3) (**Figure 1B**). There was no apparent bottleneck in the S2 and S3 screens (**Figure 1C**).

**Figure 1.**
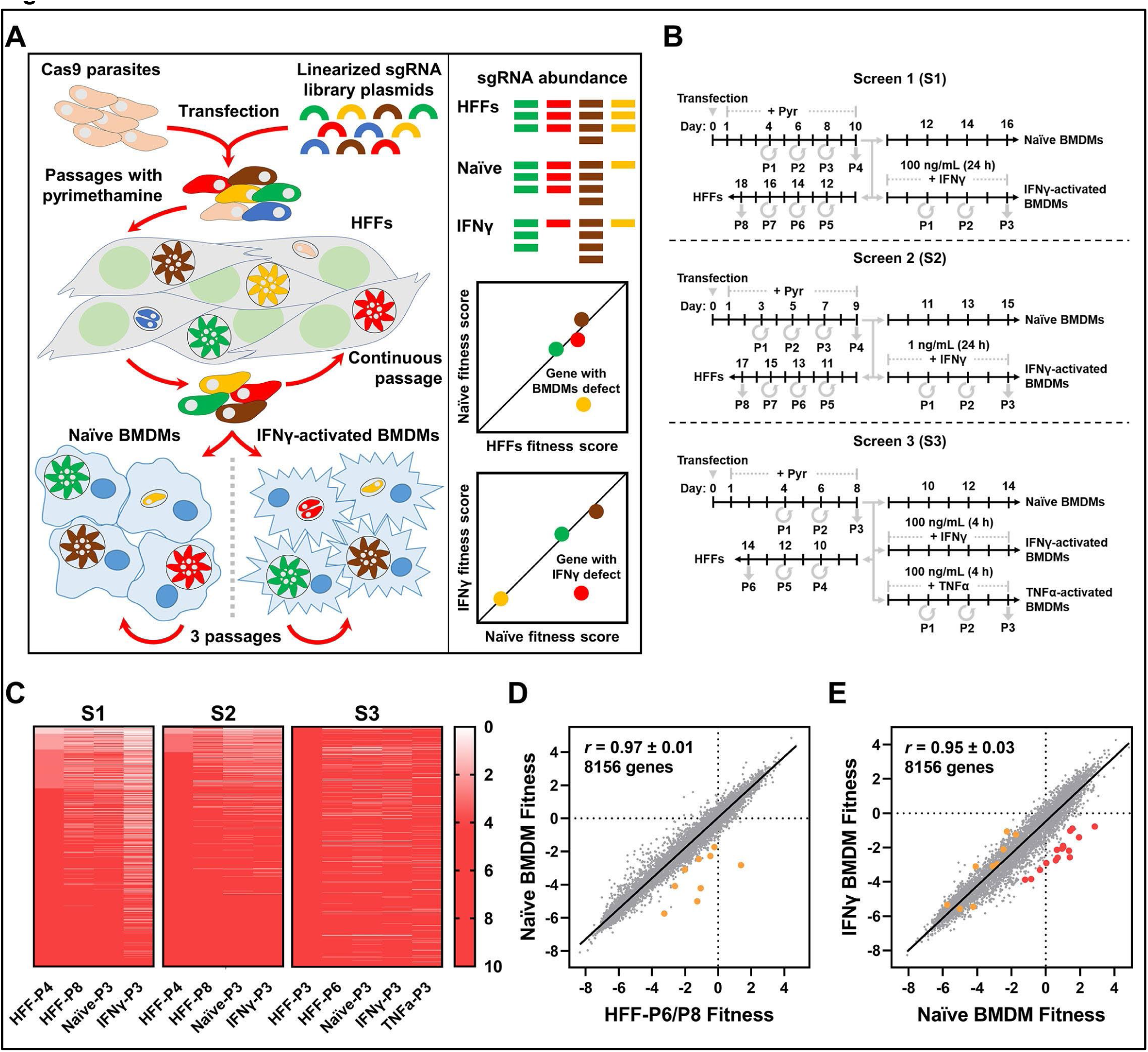
*Toxoplasma gondii* genome-wide loss-of-function screen in naïve or IFNγ-stimulated murine bone marrow-derived macrophages. (A) Screening workflow. At least 5×10^8^ Cas9-expressing RH parasites were transfected with linearized plasmids containing 10 sgRNAs against every *Toxoplasma* gene. Transfected parasites were passaged in HFFs under pyrimethamine selection to remove non-transfected parasites and parasites that integrated plasmids with sgRNAs targeting parasite genes important for fitness in HFFs. Subsequently, the pool of mutant parasites was either continuously passaged in HFFs or passaged for three rounds in murine BMDMs that were left unstimulated or pre-stimulated with IFNγ. The sgRNA abundance at different passages, determined by illumina sequencing, was used for calculating fitness scores and identify genes that confer fitness in naïve or IFNγ-activated BMDM. (B) Timeline for the generation of mutant populations and subsequent selection in the presence or absence of murine IFNγ or TNFα. Times at which parasites were passaged (P) are indicated. (C) Potential bottlenecks were assessed by determining the number of sgRNAs targeting control genes. The genes are ranked according to the number of sgRNAs in HFFs (P3 or P4) with the genes on top having the lowest sgRNA number. The scale bar indicates the sgRNA number of genes colored with graded red. Genes with 4 or more sgRNAs are colored red; otherwise, the gene is colored in a graded red scale based on how many sgRNAs were present in each sample. (D) Correlation between mean Naïve BMDM fitness and mean HFF-P6/P8 fitness. Orange dots indicate the top 9 candidate genes that confer fitness in naïve BMDMs compared to HFFs (**Table 1**). (E) Correlation between mean IFNγ-activated BMDM fitness and mean Naïve BMDM fitness of all the genes. Red dots indicate the top 16 IFNγ fitness-conferring candidate genes in murine BMDMs and orange dots the top 9 genes conferring fitness in naïve BMDMs compared to HFFs (**Table 1**).

**Table 1.** Toxoplasma genes that determine fitness in naïve or IFNγ-activated macrophages

| **Genes with a fitness defect in naïve macrophages** |  |  |  |  |  |  |
| --- | --- | --- | --- | --- | --- | --- |
| Gene ID | Product Description | Naïve BMDM vs. HFF-P6/P8 Fitness | Negative Enriched P-Value | HFF-P3/P4 Fitness | Naïve BMDM Fitness | IFNγ BMDM vs. Naïve BMDM Fitness / P-Value |
| TGGT1_249580 | *Tg*ApiAT6-3 | -4.2 | 7E-07 | 1.6 | -2.8 | -0.2 / 0.6 |
| TGGT1_253780 | Putative GTP cyclohydrolase I | -3.7 | 0.06 | 1.3 | -5.0 | -0.6 / 0.5 |
| TGGT1_247360 | PAP2 superfamily protein | -3.2 | 0.07 | 0.5 | -4.2 | -1.2 / 0.2 |
| TGGT1_305800 | 6-pyruvoyl tetrahydrobiopterin synthase | -2.5 | 0.01 | -0.3 | -5.7 | 0.4 / 0.8 |
| TGGT1_250880 | Kinase, pfkB family protein (Adenosine Kinase) | -1.8 | 0.07 | 0.8 | -2.3 | 1.2 / 0.7 |
| TGGT1_290860 | *Tg*ApiAT6-2 | -1.5 | 0.02 | 0.6 | -1.7 | 0.5 / 0.8 |
| TGGT1_243970 | Lipovitellin-like (Swiss-Model/HHpred) | -1.5 | 0.1 | -1.4 | -4.1 | 1.0 / 0.8 |
| TGGT1_224840 | 3'5'-cyclic nucleotide phosphodiesterase domain-containing protein | -1.3 | 0.04 | -0.9 | -2.5 | 0.3 / 0.6 |
| TGGT1_310000 | G-protein-coupled receptor-like (Swiss-Model/HHpred) | -1.1 | 0.09 | -0.1 | -3.1 | -7.9E-03 / 0.4 |
| **Genes with a fitness defect in IFNγ-activated macrophages** |  |  |  |  |  |  |
| Gene ID | Product Description | IFNγ BMDM vs. Naïve BMDM Fitness | Negative Enriched P-Value | HFF-P3/P4 Fitness | Naïve BMDM vs. HFF-P6/P8 Fitness | IFNγ BMDM vs. TNFα BMDM Fitness (S3) / P-Value |
| TGGT1_269620 | DNA repair protein-like (Swiss-model) | -4.0 | 0.005 | 1.0 | 1.5 | -3.0 / 0.005 |
| TGGT1_209500 | DNA repair protein-like (HHpred) | -3.6 | 2E-05 | 2.1 | 0.4 | -2.3 / 5E-05 |
| TGGT1_295340 | UV excision repair protein Rad23 protein | -3.5 | 0.002 | 1.1 | 1.4 | -3.5 / 6E-05 |
| TGGT1_316250 | GRA45 | -3.3 | 9E-04 | 1.7 | 1.0 | -2.2 / 0.03 |
| TGGT1_232670 | Cytohesin-3-like; GEF of Arf6 (Swiss-Model) | -3.3 | 0.05 | 1.3 | 0.3 | -1.9 / 0.1 |
| TGGT1_321530 | Cathepsin L (CPL) | -3.3 | 0.009 | 1.4 | 1.7 | -1.6 / 0.005 |
| TGGT1_263560 | Putative GRA<sup>#</sup>, ER-resident protein (ToxoDB) | -3.0 | 5E-04 | 1.3 | 1.5 | -1.8 / 2E-04 |
| TGGT1_205350 | Putative integral membrane protein, GNS1/SUR4 family protein | -3.0 | 0.04 | 1.2 | 0.7 | -3.7 / 5E-04 |
| TGGT1_208370 | GRA46 | -3.0 | 0.01 | 1.2 | 0.5 | -0.9 / 0.007 |
| TGGT1_231210 | Sarcalumenin/eps15 family protein (Dynamin like protein; Swiss-Model/HHpred) | -3.0 | 0.03 | 0.3 | 1.0 | -1.9 / 0.005 |
| TGGT1_228300 | CCDC25 protein | -2.9 | 0.05 | 0.8 | 0.3 | -2.3 / 0.003 |
| TGGT1_215220 | GRA22 | -2.9 | 0.03 | 1.0 | 1.1 | -0.6 / 0.4 |
| TGGT1_314500 | SUB2 | -2.8 | 0.04 | 1.3 | 0.8 | -1.7 / 4E-04 |
| TGGT1_221170 | CAAX metallo endopeptidase | -2.7 | 0.03 | -0.2 | 1.8 | -1.2 / 0.05 |
| TGGT1_200290 | ROM1 | -2.4 | 0.01 | 1.2 | 0.2 | -1.0 / 0.002 |
| TGGT1_240980 | Cytoplasmic dynein-like (Swiss-Model/HHpred) | -2.4 | 0.004 | 1.3 | 0.3 | -2.0 / 0.02 |
| TGGT1_269950* | Putative GRA<sup>#</sup> | -2.1 | 0.2 | -0.7 | -2.7 | -0.4 / 0.04 |

**Upper panel:** 9 top hits, selected from 130 naïve BMDM fitness-conferring genes, with a negative selection of sgRNAs (average $P < 0.1$ ) and $\geq 2$ -fold lower fitness (fitness score $\leq -1$ ) in naïve BMDMs vs. HFF-P6/P8. Genes without expression in murine macrophages (90) and with a Naïve BMDM fitness $\geq -1$ in two out of three screens were removed.
**Lower panel:** 16 top hits, selected from 466 IFNγ fitness-conferring genes, with a significant negative selection of sgRNAs (average $P < 0.05$ ), a $\geq 4$ -fold lower fitness (fitness score $\leq -2$ ) in IFNγ-activated vs. naïve BMDMs, and at least 4 sgRNAs present in the 3rd passage of naïve BMDMs. Genes without expression in murine macrophages (90) and with an average IFNγ BMDM fitness $\leq 0$ were removed.
\*, *TGGT1\_269950* was confirmed as an IFNγ fitness-conferring gene with individual knockout (**Figure 2**), although it does not fit the above criteria.

We defined the “fitness score” of a gene as the average log2-fold change abundance of sgRNAs targeting the gene in the condition (naïve or stimulated) relative to the input library. We calculated fitness scores at the 3^rd^/4^th^ (“HFF-P3/P4 fitness”) and 6^th^/8^th^ (“HFF-P6/P8 fitness”) passages in HFFs, or after three passages on naïve (“Naïve BMDM fitness”), IFNγ-stimulated (“IFNγ BMDM fitness”), or TNFα-stimulated (“TNFα BMDM fitness”) BMDMs (**Table S2**). The HFF-P3/P4 fitness scores were highly correlated (*r* = 0.81±0.05, mean±SD, *n*= 3) with previously determined HFFs fitness scores (47), highlighting the reproducibility of these screens and demonstrating that up to the point of infecting the BMDMs, the mutant pool was behaving as expected.

Because the intracellular BMDM and HFF environments are likely quite different, certain *Toxoplasma* genes could be important for fitness in naïve BMDMs and not HFFs. To identify this set of genes, we subtracted the HFF-P6/P8 fitness scores from the Naïve BMDM fitness scores and focussed on genes with a ≥ 2-fold fitness difference. We also calculated the negative enrichment of sgRNAs between Naïve BMDM 3^rd^ passage (P3) and HFF-P6/P8 by using the Model-based Analysis of Genome-wide CRISPR/Cas9 Knockout (MAGeCK) algorithm (50). We identified ∼130 parasite genes, including 9 top hits (**Figure 1D and Table 1**), with ≥ 2-fold lower fitness and/or a significant negative enrichment of sgRNAs (p < 0.05) in Naïve BMDM P3 *vs.* HFF-P6/P8 in at least two out of the three screens (and ≥ 3 sgRNAs present in HFF-P6/P8) (**Table S2**). To determine if these *Toxoplasma* genes shared biological annotation we performed gene set enrichment analysis (GSEA) (51). We found significant enrichment of *Toxoplasma* genes involved in folate metabolism (TGGT1_305800, TGGT1_253780, TGGT1_231920, TGGT1_212740, TGGT1_266366 (BT1 folate transporter (52)), purine metabolism (TGGT1_224840, TGGT1_257080, TGGT1_250880 (adenosine kinase), TGGT1_243580, TGGT1_282190), parasite membrane-localized transporters (TGGT1_226110, TGGT1_315560 (*Tg*ABCG77), *Tg*ApiAT3-1, *Tg*ApiAT6-2, *Tg*ApiAT6-3) and genes encoding for proteins with a Surface Antigen 1 Related Sequence (SRS) domain, which are glycosylphosphatidyl (GPI)-linked developmentally-regulated surface proteins that are involved in attachment/invasion (53) (**Table S3**). Overall these data suggest that the nutrient availability to *Toxoplasma* in naïve BMDMs and HFFs is variable and that *Toxoplasma* uses different SRS proteins to attach/invade BMDMs *vs*. HFFs.

Next, we subtracted the Naïve BMDM fitness scores from the IFNγ BMDM fitness scores to identify *Toxoplasma* genes that specifically determine fitness in IFNγ-stimulated BMDMs. In addition, we identified genes with a negative enrichment of sgRNAs between naïve and IFNγ-stimulated BMDMs at P3 (**Table S2**). We identified ∼466 parasite genes with ≥ 2-fold lower fitness and/or a significant negative enrichment of sgRNAs (p < 0.05) in IFNγ-stimulated, relative to naïve BMDMs P3 in at least two out of the three screens (and ≥ 3 sgRNAs present in naïve BMDMa) (**Table S2**). ROP18 and GRA12, which are known to determine RH resistance to IFNγ, had a 2.2 fold and 5.2 fold lower fitness in IFNγ-stimulated BMDMs, respectively. It should be noted that ROP5 was not in our list of hits because sgRNAs against *ROP5* were not in our library because the sgRNA design algorithm excluded *ROP5*-targeting sgRNAs because no uniquely targeting sgRNAs could be designed as *ROP5* is a multi-copy gene. Genes encoding for proteins involved in DNA repair (TGGT1_206400, TGGT1_220120, TGGT1_251670, TGGT1_272900, TGGT1_295340, TGGT1_307650, TGGT1_320110), nucleic acid binding proteins - especially proteins containing RNA-binding/recognition motifs - were significantly enriched (**Table S3**). In addition, we performed pathway analysis on the RNAseq data of activated macrophages to identify pathways that are uniquely regulated by IFNγ. The cholesterol biosynthesis pathway (**Figure S1D and S1E**) and many other metabolic pathways were significantly downregulated in IFNγ-stimulated but not in TNFα-stimulated BMDMs (**Table S1**). Indeed, most of the *Toxoplasma* genes that determine fitness in IFNγ-stimulated BMDM and were involved in lipid biosynthesis/transport had fitness defects specifically in IFNγ-stimulated cells (**Table S1**), suggesting that they are required for responding to changes in cholesterol levels in IFNγ-stimulated BMDMs. For example, genes involved in lipid biosynthesis or transport were enriched in IFNγ-activated BMDMs: three GNS1/SUR4 family members involved in long-chain fatty acid elongation and sphingolipid/ceramide biosynthesis (TGGT1_253880, TGGT1_205350, TGGT1_242380), TGGT1_229140 (enoyl-CoA hydratase like-beta-oxidation), GRA38 (an orthologue of GRA39 of which the knockout has been shown to accumulate lipid (54)), ATP-binding cassette (ABC) transporters ABCG87 and ABCG96, which have been shown to mediate cholesterol transport (55), and TGGT1_266640 (Acetyl-coenzyme A synthetase 2). A GNS1/SUR4 family member, TGGT1_205350, was recently identified as a gene important during *in vivo* infection of the peritoneum (49). IFNγ stimulation significantly downregulated the cholesterol biosynthesis pathway in BMDMs as has been shown by others (56). It is, therefore, possible that *Toxoplasma* needs to modify its acquisition/export or own production of lipids in IFNγ-stimulated BMDMs. The top 16 candidate genes that differed in fitness in IFNγ-stimulated *vs*. naïve BMDM at P3 and were specifically depleted in IFNγ-activated BMDMs (**Figure 1E**) are presented in **Table 1**. This table contains the ROM1 and SUB2 proteases, important for microneme proteins (MICs) processing and rhoptry protein maturation, respectively (57, 58), possibly indicating the role of specific MICs and ROPs in the invasion of and survival in IFNγ-activated macrophages; *Tg*CPL, important for the digestion of host cytosolic proteins in the parasite lysosomal-like organelle called the VAC (59); TGGT1_221170, a putative CAAX Metallo endopeptidase involved in the cleavage of isoprenylated proteins; three putative DNA repair proteins TGGT1_269620, TGGT1_209500, and TGGT1_295340; three proteins involved in intracellular trafficking, TGGT1_232670 (putative Cytohesin-3), TGGT1*_*231210 (putative dynamin-like protein), and TGGT1*_*240980 (putative cytoplasmic dynein); and four dense granule proteins GRA22, GRA45, GRA46, and a putative GRA, TGGT1*_*263560.

Taken together, we have identified multiple *Toxoplasma* genes that appear to confer fitness specifically in naïve or IFNγ-stimulated BMDM, likely because they are needed to adapt to differences in levels of nutrients *Toxoplasma* acquires from its host cells and to the multiple other pathways that are modulated by IFNγ.

### Single gene knockouts from hits identified by CRISPR screens have fitness defects in IFNγ-activated BMDM

To confirm some of the parasite genes that determine fitness in IFNγ-activated BMDMs, we generated individual knockouts of four genes (*TGGT1_269620*, *GRA45*, *TGGT1_232670,* and *TGGT1_263560*) in luciferase-expressing RH parasites (**Figure S2A to S2F**). All knockouts had reduced growth compared to wild-type parasites in IFNγ-activated BMDMs as determined by luciferase assays at 24 h p.i. (**Figure 2A to 2D**). We previously determined the contribution to the *in vivo* fitness of 217 *Toxoplasma* genes (49). Of the 24 genes that determined *in vivo* fitness, two genes (*TGGT1_205350* and *GRA22*) were present in the top hits for fitness in IFNγ-activated BMDMs (**Table 1**). We previously confirmed that Δ*gra22* and Δ*TGGT1*_*269950* parasites were outcompeted by wild-type in the peritoneum. IFNγ-stimulated BMDMs significantly inhibited Δ*gra22* and Δ*TGGT1*_*269950* parasite growth relative to wild-type parasites (**Figure 2E, 2F**). These results indicate that the *in vivo* fitness defect of these two genes is likely due to their importance in determining parasite fitness in IFNγ-stimulated peritoneal macrophages. Most of the knockouts had no, or a minor, fitness defect in mouse embryonic fibroblasts (MEFs) or naïve murine BMDMs (**Figure S3A to S3G**). Δ*TGGT1_263560* and Δ*gra22* parasites formed larger plaques compared to wild-type parasites in MEFs, although this was only significant for Δ*TGGT1_263560* (**Figure S3G**). Although the Δ*TGGT1_263560* plaques were larger, they appeared quite different compared to wild-type plaques; the lysis area contained very few parasites and host cells, possibly due to an early egress phenotype (**Figure S3H**). To test the impact of candidate gene knockout in a non-luciferase expressing type 1 (RH) strain, we performed a growth competition assay, in which growth differences between wild-type and knockouts accumulate over three passages in both naïve and IFNγ-activated BMDMs. As a control, Δ*gra22* parasites were outcompeted by wild-type parasites only in the IFNγ-activated but not naïve BMDMs (**Figure 2G**). Similar to Δ*gra22* parasites, parasites lacking *ROM1* (**Figure S2G**) were outcompeted by wild-type parasites in IFNγ-activated BMDMs (**Figure 2H**). Thus, the parasite genes identified by our genome-wide loss-of-function screen indeed determine parasite fitness in IFNγ-activated BMDM.

**Figure 2.**
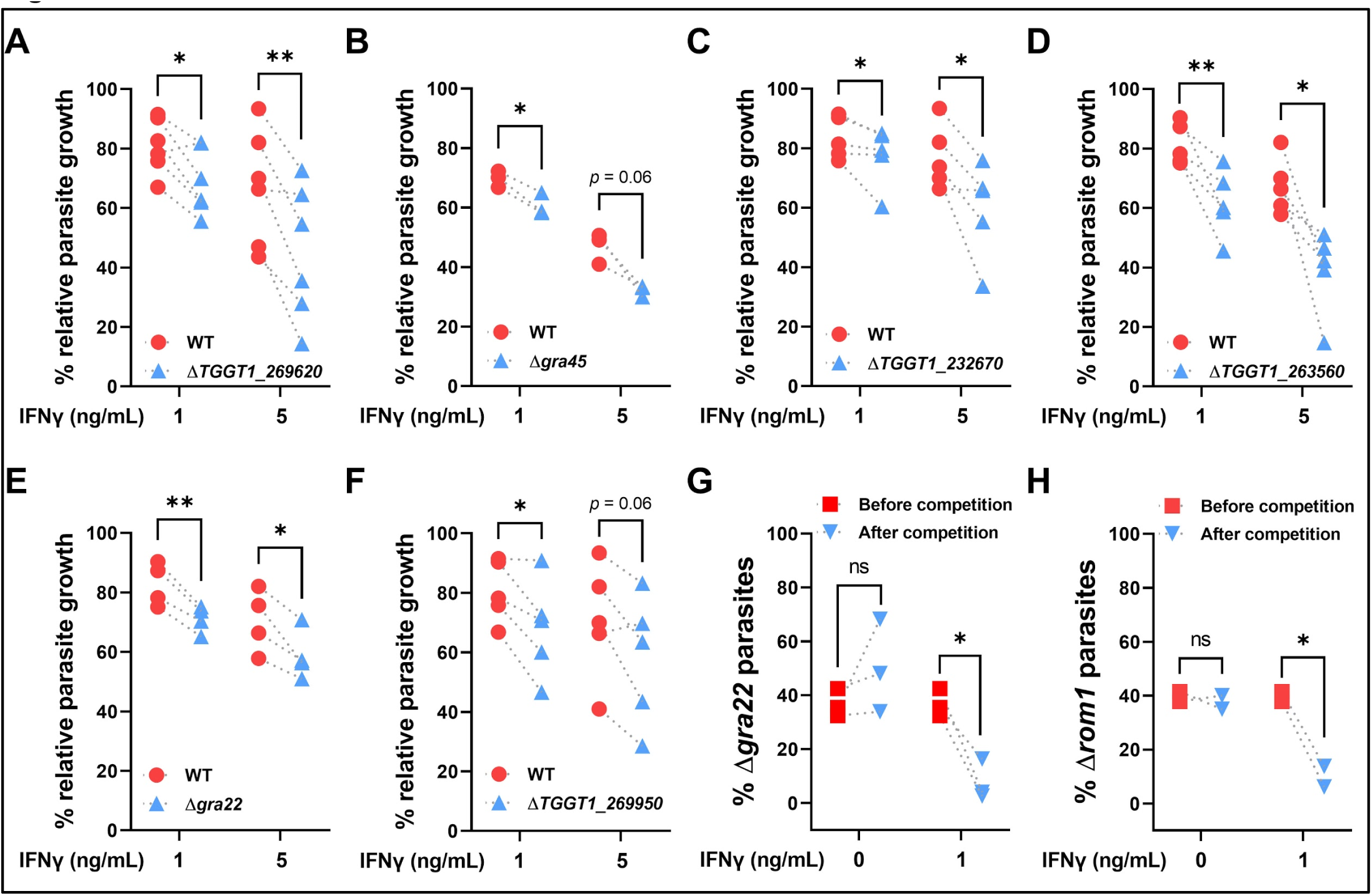
Validation of candidate genes that determine parasite fitness in IFNγ-stimulated murine BMDMs. (A to F) Murine BMDMs pre-stimulated with IFNγ (1 or 5 ng/mL) or left unstimulated for 24 h were infected with luciferase-expressing wild-type (WT) parasites or with parasites in which (A) *TGGT1_269620*, (B) *GRA45*, (C) *TGGT1_232670*, (D) *TGGT1_263560*, (E) *GRA22* or (F) *TGGT1_269950* was knocked-out (MOI of 0.25). 24 h p.i. parasite growth for each strain was measured by luciferase assay. Parasite growth in IFNγ-activated BMDMs is expressed relative to growth in naïve BMDMs. Data are displayed as paired plots of at least 3 independent experiments. Asterisk (*) indicates a significant difference between WT and knockout analyzed with two-tailed paired *t* test: (A) *n* = 6, *p* = 0.02 for 1 ng/mL IFNγ, *p* = 0.007 for 5 ng/mL IFNγ; (B) *n* = 3, *p* = 0.02 for 1 ng/mL IFNγ; (C) *n* = 5, *p* = 0.04 for 5 ng/mL IFNγ; (D) *n* = 5, *p* = 0.006 for 1 ng/mL IFNγ, *p* = 0.01 for 5 ng/mL IFNγ; (E) *n* = 4, *p* = 0.006 for 1 ng/mL IFNγ, *p* = 0.02 for 5 ng/mL IFNγ; (F) *n* = 5, *p* = 0.03 for 1 ng/mL IFNγ. (G and H) Growth competition assay between GFP-positive WT parasites and GFP-negative (G) Δ*gra22* or (H) Δ*rom1* parasites was performed in murine BMDMs pre-stimulated with 1 ng/mL IFNγ or left unstimulated for 3 passages. The percentage of Δ*gra22* or Δ*rom1* was determined at the start of the competition and after 3 passages by plaque assay measuring the GFP-negative plaques *vs*. total plaques. Data are displayed as paired plots for Δ*gra2*2 (*n* = 3) and for Δ*rom1* (*n* = 2). Asterisk (*) indicates a significant difference between the ratio of WT *vs*. knockout before and after competition analyzed with two-tailed paired *t* test: (G) *p* = 0.04 for 1 ng/mL IFNγ-activated BMDMs; (H) *p* = 0.04 for 1 ng/mL IFNγ-activated BMDMs.

### GRA45, a top hit from the screen, is important for parasite fitness in IFNγ-stimulated macrophages of other species

Of the seven genes we confirmed to have a fitness defect in IFNγ-activated BMDM, four genes (*GRA22*, *GRA45*, *TGGT1_263560*, and *TGGT1_269950*) encode for known or putative GRAs (45, 54, 60). However, of these four GRAs, only GRA22 was previously characterized and shown to contribute to parasite natural egress (60), while GRA45 was recently discovered as an ASP5 substrate with unknown function (45). Macrophages from different species have different mechanisms to control *Toxoplasma* growth (8) and therefore knowing if these genes are important for conferring resistance to IFNγ in other species could help us narrow down the possible mechanism by which they do so. To test if these two GRAs are also important for parasite fitness in IFNγ-stimulated macrophages of other species, primary rat BMDMs and human THP-1 macrophages were stimulated with IFNγ or left unstimulated followed by infection with wild-type, Δ*gra22* or Δ*gra45* parasites (**Figure 3**). Δ*gra45* parasites, but not Δ*gra22* parasites, were significantly more susceptible to IFNγ-mediated growth inhibition compared to wild-type parasites in both rat and human macrophages (**Figure 3**) indicating GRA45 plays an important function in the resistance of *Toxoplasma* to IFNγ in multiple species. We therefore decided to focus on determining the function of GRA45.

**Figure 3.**
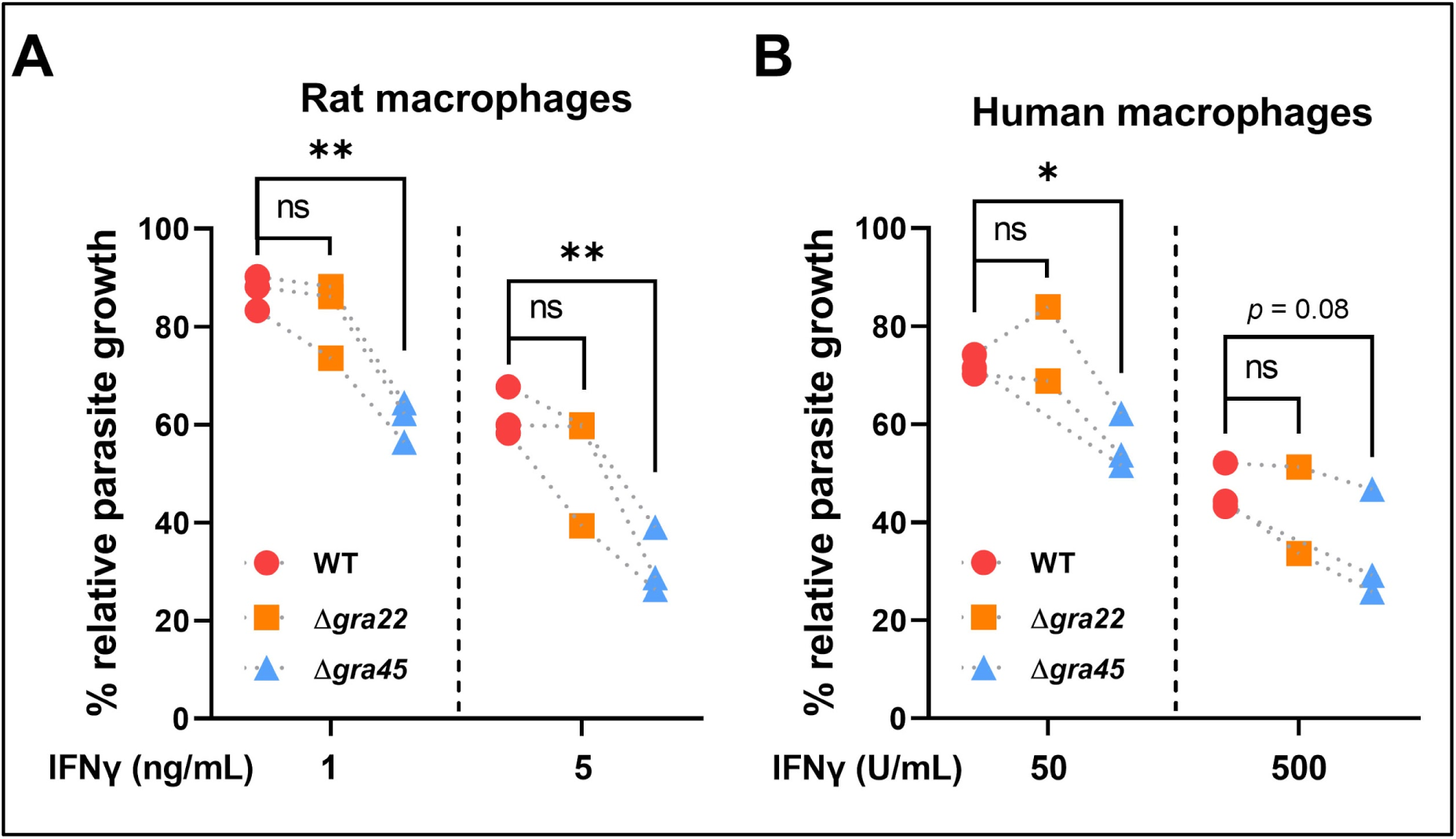
Δ*gra45* parasites, but not Δ*gra22* parasites, have enhanced susceptibility to IFNγ-mediated growth inhibition in rat BMDMs and human THP-1 macrophages. (A) Brown Norway rat BMDMs pre-stimulated with or without IFNγ (1 ng/mL or 5 ng/mL) were infected with luciferase-expressing WT, Δ*gra22* or Δ*gra45* parasites (MOI of 0.25) for 24 h. Parasite growth for each strain was measured by luciferase assay and the growth in IFNγ-activated BMDMs is expressed relative to growth in naïve BMDMs. Data are displayed as paired plots for 3 independent experiments. Asterisk (*) indicates a significant difference analyzed by one-way ANOVA with Tukey’s multiple comparisons test: *p* = 0.007 for WT *vs.* Δ*gra45* in BMDMs stimulated with 1 ng/mL IFNγ, *p* = 0.002 for WT *vs.* Δ*gra45* in BMDMs stimulated with 5 ng/mL IFNγ. (B) PMA-differentiated THP-1 macrophages pre-stimulated with or without IFNγ (50 U/mL or 500 U/mL) were infected with luciferase-expressing WT, Δ*gra22* or Δ*gra45* parasites (MOI of 0.25) for 24 h. Parasite growth for each strain was measured by luciferase assay and the growth in IFNγ-activated BMDMs is expressed relative to growth in naïve BMDMs. Data are displayed as paired plots with n = 2 for Δ*gra22* and *n* = 3 for WT and Δ*gra45.* Due to different *n* between Δ*gra22* and Δ*gra45*, the significant difference was analyzed with two-tailed paired *t* test. Asterisk (*) indicates *p* = 0.02 for WT *vs.* Δ*gra45* in THP-1 macrophages stimulated with 50 U/mL IFNγ.

### GRA45 has structural homology to protein chaperones

GRA45 was recently shown to interact with GRA44 (TGGT1_221870) and With-No-Gly-loop (WNG)2 kinase (TGGT1_240090), which also localize to dense granules. GRA45, GRA44 and WNG2 are relatively conserved and orthologues are present in coccidian species belonging to the Sarcocystidae (*Neospora caninum*, *Hammondia hammondi*, *Cystoisospora suis*, *Sarcocystis neurona*) and Eimeriidae (*Eimeria* spp. and *Cyclospora cayetanensis*). This is a rather unique phylogenetic profile as most GRAs are only conserved within the *Toxoplasmatinae* suggesting that these GRAs have a conserved function in these different parasite species. GRA45 is highly expressed in tachyzoite and bradyzoite stages and has moderate expression levels in sexual stages (ToxoDB.org). A paralogue of GRA45 exists in *Toxoplasma* (TGGT1_295390), which is exclusively expressed in sexual stages. Interestingly, GRA44 is duplicated and one copy (TGGT1_221860) is also exclusively expressed in sexual stages. Searches for primary sequence homology or known protein domains failed to provide any suggestions for putative GRA45 functions. However, homology searches based on secondary structure prediction indicated that the N-terminal region of GRA45 (from amino acids 72 to 225) has structural homology to small heat shock proteins (sHSPs) (**Figure S4A**), which are ATP-independent chaperones that prevent protein aggregation. The conserved [I/V/L]x[I/V/L] signature present at the C-terminus of most sHSP was conserved in GRA45 (RID<u>V</u>H from amino acid 204 to 208) and its orthologues. The most significant structural homology was to Aggregation Suppressing Protein A (AspA) from *Salmonella typhimurium* (61) and HSP17.9 from *Xylella fastidiosa* (62). The GRA45 C-terminal region (amino acids 260-335) had structural homology to DUF1812 domains (63) and Mfa2 (64), which share a transthyretin-like fold (**Figure S4A**). Transthyretin family members are known to transport lipids such as retinol and thyroxine (65). In addition, the top 10 models with GRA45 structural homology, as predicted by I-TASSER (66), are predicted to bind hydrophobic substrates as they have acyltransferase domains. Overall these data indicate that a potential function of GRA45 is binding of hydrophobic substrates and preventing their aggregation. GRA45 possesses an N-terminal TEXEL (RRL) motif that is cleaved by ASP5 (45). Consistent with published results that GRA45 remains in the PV lumen after secretion (45), we observed that almost all GRA45 co-localized with PV-resident GRA1 and GRA2 in the PV lumen but only partially with PVM-associated GRA5 and GRA7, even though these GRAs all localize to dense granules in extracellular parasites (**Figure S4B**). Thus, GRA45 is unlikely to be a *Toxoplasma* effector that directly modulates or inhibits IFNγ-induced host effector mechanisms.

### Deletion of GRA45 leads to aggregation of dense granule proteins inside the parasite

Based on its structural homology to sHSPs, we hypothesized that instead of directly neutralizing IFNγ-mediated inhibition GRA45 might perform a chaperone-like function by binding to the hydrophobic domain present in many GRA effectors, thereby preventing their aggregation. GRAs with a transmembrane domain have peculiar trafficking and do not traffic to the parasite plasma membrane but instead traffic to the dense granules where they exist in a soluble state (67). What prevents their transmembrane domain from inserting into the ER membrane or the dense granule membrane is unknown but has been hypothesized to be mediated by a dense-granule-localized chaperone (67, 68). After exocytosis of the dense granules, these transmembrane-containing GRAs somehow integrate as type I transmembrane proteins into the PVM and/or intravacuolar network (IVN). To test the hypothesis that GRA45 plays a role in preventing the aggregation of GRAs, we examined the solubility of hydrophobic GRAs, such as GRA2, which has two amphipathic helices that mediate its membrane association post-secretion (69), and GRA7, which contains a transmembrane domain but localizes to the dense granule core and does not integrate into the dense granule membrane (70, 71). In wild-type parasite lysates generated by freeze/thawing, GRA2 and GRA7 were mainly present in the pellet fraction and ran according to their corresponding molecular weight (MW) in denaturing conditions (**Figure 4A, left panels**). In addition to their corresponding MW, in Δ*gra45* parasites, GRA2 and GRA7 appeared at a much higher MW as bands that were not completely separated during the gel migration consistent with protein aggregation. These high MW bands of GRA2 and GRA7 were barely observed in wild-type or in Δ*gra45* parasites complemented with an HA-tagged version of *GRA45* (Δ*gra45*+*GRA45HA*, **Figure S2C**) (**Figure 4A, left panels**). As a control, we used the parasite plasma membrane protein SAG1, which traffics to the parasite surface independent of dense granules and should therefore not be affected by GRA45. No higher MW bands of SAG1 were observed in any of the parasite strains (**Figure 4A, left panels**) indicating that the aggregation of GRA2 or GRA7 was not due to excess sample loading. After treatment with non-ionic detergents (NP-40 or Triton X-100), which solubilize membrane proteins by associating with their hydrophobic surface, non-aggregated GRA2 and GRA7 running at the expected MW were completely solubilized and released into the supernatant fraction in all the strains. However, in the Δ*gra45* parasites, the high MW bands of GRA2 and GRA7 were still mostly present in the pellet fraction (**Figure 4A, middle and right panels**). The presence and the persistence of high MW bands corresponding to GRA2 and GRA7 in Δ*gra45* parasites are consistent with their aggregation, possibly *via* noncovalent hydrophobic interactions inside the dense granules.

**Figure 4.**
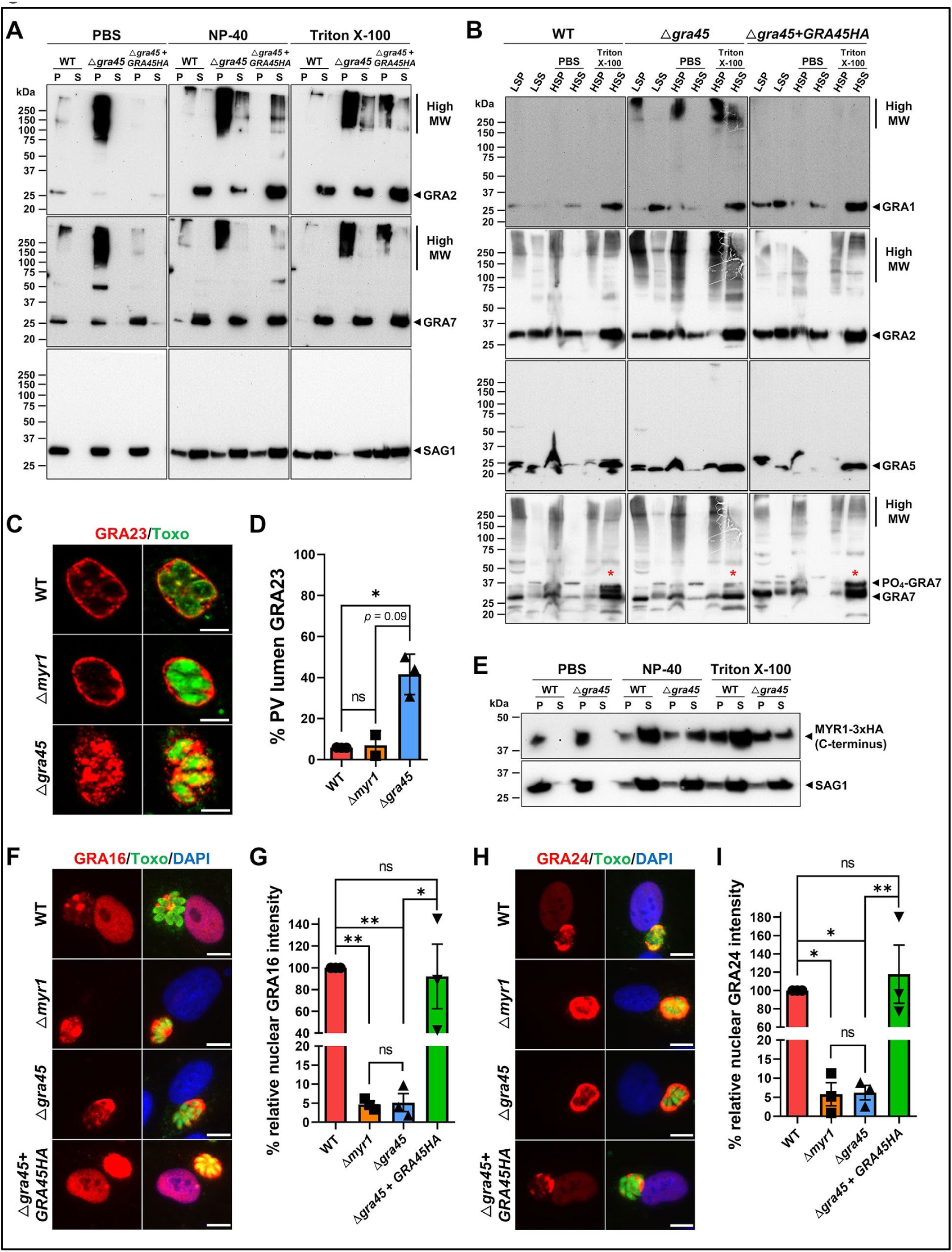
In Δ*gra45* parasites other GRAs are aggregated and mislocalized after secretion into the vacuole. (A) Extracellular parasites of WT, Δ*gra45* or Δ*gra45* + *GRA45HA* strains were disrupted by freeze/thaw cycles followed by incubating with PBS, 1% NP-40 or 1% Triton X-100 and fractionating by high-speed centrifugation to separate the pellet (P) and supernatant (S). GRA2 and GRA7 were detected with indicated antibodies. SAG1 was used as the parasite loading control. The image is representative of results from two independent experiments. (B) HFFs were infected with WT, Δ*gra45*, or Δ*gra45* + *GRA45HA* parasites for 24 h followed by mechanical disruption of host cells and separating intact parasites (LSP) and membrane fraction (LSS) *via* low-speed centrifugation. The PVM fraction was harvested and incubated with PBS, 1% NP-40 or 1% Triton X-100 and fractionated by high-speed centrifugation to separate the pellet (HSP) and supernatant (HSS). Western blots were developed with antibodies against GRA1, GRA2, GRA5, and GRA7. Red stars indicate the samples containing a phosphorylated form of GRA7. (C) HFFs were infected with WT, Δ*gra45* or Δ*myr1* parasites that transiently expressed GRA23-HA followed by fixing and staining with antibodies against SAG1 (green) and the HA epitope (red). The images are representative of results from two independent experiments and were taken at identical exposure times for each channel (scale bar = 5 μm). (D) Localization of GRA23 in at least 50 vacuoles containing 4 or more parasites was observed and the percentage of PV lumen localized GRA23 was quantified. Data are displayed as averages ± SD with n = 2 for Δ*myr1* and *n* = 3 for WT and Δ*gra45.* Due to different *n* between Δ*myr1* and Δ*gra45*, the significant difference was analyzed with two-tailed paired *t* test. Asterisk (*) indicates a *p* = 0.02 for WT *vs.* Δ*gra45*. (E) Extracellular WT parasites expressing endogenously 3xHA-tagged MYR1 or Δ*gra45* parasites in this background were disrupted by freeze/thaw cycles followed by incubating with PBS, 1% NP-40 or 1% Triton X-100 and fractionated by high-speed centrifugation to separate the pellet (P) and supernatant (S). The C-terminal polypeptide of MYR1 was detected with anti-HA antibodies. SAG1 was used as the parasite loading control. (F) WT, Δ*myr1*, Δ*gra45*, or Δ*gra45* + *GRA45HA* parasites were transiently transformed with GRA16-HA-FLAG expressing plasmid and immediately used to infect HFFs and fixed at 24 h p.i. and subjected to the immunofluorescent assay with indicated antibodies. The image is representative of results from three independent experiments and was taken at identical exposure times for each channel (scale bar = 10 μm). (G) GRA16 nuclear intensity was quantified in host cells containing a single parasitophorous vacuole with 4 or more parasites for at least 30 host cells. Data are displayed as averages ± SD relative to GRA16 nuclear intensity of WT-infected cells for 3 independent experiments. Asterisk (*) indicates a significant difference analyzed with one-way ANOVA with Tukey’s multiple comparisons test: *p* = 0.008 for WT *vs.* Δ*myr1*, *p* = 0.009 for WT *vs.* Δ*gra45*, *p* = 0.01 for Δ*gra45 vs.* Δ*gra45* + *GRA45HA*. (H) WT, Δ*myr1*, Δ*gra45*, or Δ*gra45* + *GRA45HA* parasites were transiently transformed with a GRA24-HA-FLAG expressing plasmid and immediately used to infect HFFs and fixed at 24 h p.i. and subjected to the immunofluorescent assay with indicated antibodies. The image is representative of results from three independent experiments and was taken at identical exposure times for each channel (scale bar = 10 μm). (I) GRA24 nuclear intensity was quantified in host cells containing a single parasitophorous vacuole with 4 or more parasites for at least 30 host cells. Data are displayed as averages ±SD relative to GRA24 nuclear intensity of WT-infected cells for 3 independent experiments. Asterisk (*) indicates a significant difference analyzed with one-way ANOVA with Tukey’s multiple comparisons test: *p* = 0.01 for WT *vs.* Δ*myr1*, *p* = 0.01 for WT *vs.* Δ*gra45*, *p* = 0.005 for Δ*gra45 vs.* Δ*gra45* + *GRA45HA*.

To determine if the solubility of GRAs was also affected after exocytosis into the PV we performed cell fractionation of intracellular parasites. Intact parasites, mechanically released from host cells and pelleted and resuspended in lysis buffer containing DTT, are referred to as the low-speed pellet (LSP). The supernatant (LSS), which contains PVM and host cell material, was fractionated with high-speed centrifugation to isolate the supernatant (HSS) and pellet (HSP) fractions. The HSP-containing PVM and other membrane fractions (for example, the IVN) was subjected to detergent treatment to extract the membrane-associated GRAs or left untreated. We examined different GRAs based on their final destinations in the PV and their membrane association properties: GRA1, localizes to the PV lumen as a soluble protein (72, 73); GRA2, is an IVN-associated protein (72); GRA5, is a PVM-integrated protein that faces to the host cytosol (74, 75); and GRA7, associates with IVN and PVM (70, 71), and strands extending from PVM into the host cytosol (76). In all strains, GRA1 is mainly found in the soluble fraction and also in the LSP that contains intact parasites. However, in Δ*gra45* parasites the LSP of intact parasites and the insoluble HSP fraction without detergent treatment contained high MW GRA1, while it remained in both Triton X-100-treated HSP and HSS (**Figure 4B**). The insoluble GRA2 was extracted from the HSP after Triton X-100 treatment in all parasite strains, but the high MW of GRA2 was resistant to extraction by Triton X-100 in Δ*gra45* parasites (**Figure 4B**). Although we did not observe GRA5 high MW bands, GRA5 in the insoluble fraction was less extracted from the HSP after Triton X-100 treatment in Δ*gra45* parasites compared to wild-type and Δ*gra45*+*GRA45HA* parasites (**Figure 4B**). Intracellularly, GRA7 is present as a non-phosphorylated protein running at 26 kDa and a phosphorylated protein running at 32 kDa (**Figure 4B**), which is only present in infected host cells in the PVM where it faces to the host cytosol (77). GRA7 was also well extracted from the HSP after Triton X-100 treatment in all the parasite strains, but in Δ*gra45* parasites the phosphorylated form of GRA7 was less abundant compared to wild-type and Δ*gra45*+*GRA45HA* parasites (**Figure 4B**) indicating GRA7 was not properly localized to the PVM in Δ*gra45* parasites. Taken together, these results indicate that without GRA45 the GRAs inside the dense granules aggregate, which subsequently appears to lead to their mislocalization after exocytosis into the PV.

### Deletion of GRA45 leads to the mislocalization of GRAs upon exocytosis of dense granules

Because we observed aggregation of GRA proteins in the parasite we wanted to determine if the localization of GRAs after exocytosis of the dense granules was affected. We initially focused on GRA23, as it is almost exclusively localized to the PVM (78, 79), where it plays a role as a molecular sieve (80) allowing passive size-dependent import of host-derived small molecules. However, in Δ*gra45* parasites, a large fraction of GRA23 was mislocalized to the PV lumen whereas the localization remained normal in the parasites lacking MYR1 (Δ*myr1*) (**Figure S2D**), which is served as a control as it was shown that Δ*myr1* parasites have normal function of PVM-localized GRAs (42) (**Figure 4D**). Given that GRA7 and GRA23, which are normally localized to the PVM, appeared to be mislocalized in Δ*gra45* parasites, we hypothesized that components of the translocon involved in GRA translocation across the PVM could also be mislocalized. To test this hypothesis, we generated Δ*gra45* parasites in a strain where MYR1 was endogenously tagged with 3xHA epitopes (**Figure S2C and S2I**). Although we did not observe any high MW bands corresponding to MYR1 in Δ*gra45* parasites (**data not shown**), MYR1 was more resistant to detergent extraction and less abundant in the soluble fraction of Δ*gra45* parasites compared to wild-type parasites (**Figure 4E**) indicating GRA45 is also involved in maintaining MYR1 soluble. However, it is unclear what happens to the N-terminus of MYR1 in Δ*gra45* parasites, as MYR1 is cleaved by ASP5 into an N-terminal and C-terminal polypeptide (42, 81) and only the C-terminal MYR1 polypeptide was observed for this experiment. Furthermore, MYR2 and MYR3 are also required for GRA effector translocation into the host cell (43). We therefore indirectly assessed if a translocon-mediated phenotype was affected in Δ*gra45* parasites by determining GRA16 and GRA24 export beyond the PVM. Indeed, GRA16 and GRA24 were no longer exported beyond the PVM in Δ*gra45* parasites, as observed by their absence from host nuclei (**Figure 4F and 4H**), which was similar to the Δ*myr1* phenotype (**Figure 4G and 4I**). Overall, these data indicate that GRA45 either affects the correct assembly of the translocon into the PVM or the correct trafficking of GRA effectors to the translocon.

### GRA45 is important for parasite virulence

Since the data suggest that GRA45 affects the correct localization of GRA proteins to the PVM and export beyond the PVM and without GRA45 many GRAs aggregate, it is conceivable that the fitness defect of Δ*gra45* parasites in IFNγ-stimulated BMDMs was due to the defect of GRA effector export beyond the PVM. However, the translocon components MYR1/2/3 did not emerge as hits in our loss-of-function screen in IFNγ-stimulated BMDMs (**Figure 5A**). GRA45 has a significantly lower IFNγ BMDM *vs*. Naïve BMDM fitness score compared to MYR1/2/3 across the three screens (**Figure 5B**). To confirm the results from the screen we compared side-by-side the susceptibility of Δ*gra45* and Δ*myr1* parasites to IFNγ-mediated growth inhibition. In murine BMDMs pre-stimulated with increasing concentrations of IFNγ, we observed that Δ*gra45*, but not Δ*myr1*, parasites were more susceptible than either wild-type or Δ*gra45*+*GRA45HA* parasites to IFNγ-mediated growth inhibition (**Figure 5C**). To determine if these *in vitro* differences also translated to differences in virulence in mice, we intraperitoneally (i.p.) infected mice with 100 tachyzoites of wild-type, Δ*gra45*, Δ*myr1*, Δ*gra45+GRA45HA* parasites and monitored parasite virulence. Consistent with the *in vitro* data, Δ*gra45*, but not Δ*myr1*, parasites were significantly less virulent compared to wild-type or GRA45 complemented parasites (**Figure 5D**). These data indicate that the effect of GRA45 on parasite virulence cannot be solely explained by the defect in GRA export beyond the vacuole but is likely due to the combined defect in GRA export beyond the vacuole and the defect in correct GRA localization in the PV.

**Figure 5.**
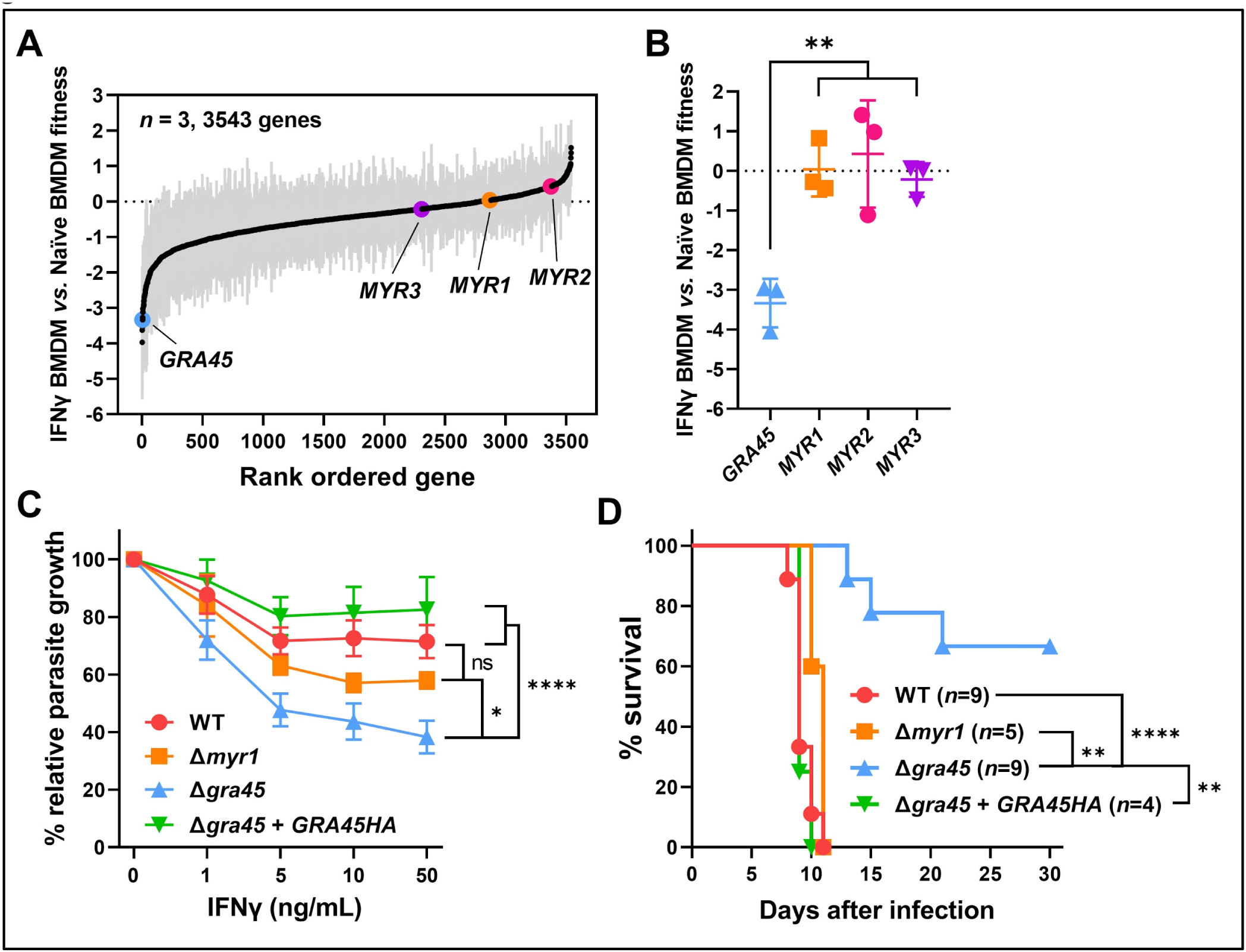
Compared to Δmyr1 parasites, Δgra45 parasites are more susceptible to IFNγ and less virulent in mice. (A) *Toxoplasma* genes that have at least 3 sgRNAs present after the 3rd passage in naïve BMDM in the three screens are rank-ordered according to their IFNγ *vs*. naïve BMDM fitness scores. Data are displayed as average fitness scores (black plots) ±SD (gray lines). (B) IFNγ *vs*. naïve BMDM fitness scores of *GRA45*, *MYR1*, *MYR2*, and *MYR3*. Data are displayed as averages ± SD for 3 independent screens. Asterisk (*) indicates a significant difference analyzed with one-way ANOVA with Tukey’s multiple comparisons test: *p* = 0.005 for *GRA45 vs. MYR1*, *p* = 0.003 for *GRA45 vs. MYR2*, *p* = 0.009 for *GRA45 vs. MYR3*. (C) Murine BMDMs pre-stimulated with or without IFNγ (1 ng/mL, 5 ng/mL, 10 ng/mL or 50 ng/mL) were infected with luciferase-expressing WT, Δ*myr1*, Δ*gra45*, or Δ*gra45* + *GRA45HA* parasites (MOI of 0.25) for 24 h. Parasite growth for each strain was measured by luciferase assay and the growth in IFNγ-activated BMDMs is expressed relative to growth in naïve BMDMs. Data are displayed as connecting lines with an average of four independent experiments ± SD. Asterisk (*) indicates a significant difference analyzed with two-way ANOVA with Tukey’s multiple comparisons test: *p* < 0.0001 for WT *vs.* Δ*gra45*, *p* < 0.0001 for Δ*gra45 vs.* Δ*gra45* + *GRA45HA*, *p* = 0.01 for Δ*myr1 vs.* Δ*gra45*. (D) CD-1 mice were i.p. infected with 100 tachyzoites of WT, Δ*myr1*, Δ*gra45* or Δ*gra45* + *GRA45HA* parasites and survival of mice was monitored for 30 days. Data are displayed as Kaplan-Meier survival curves. Asterisk (*) indicates a significant difference analyzed with Log-rank (Mantel-Cox) test: *p* < 0.0001 for WT *vs.* Δ*gra45*, *p* = 0.006 for Δ*gra45 vs.* Δ*gra45* + *GRA45HA*, *p* = 0.007 for Δ*myr1 vs.* Δ*gra45*.

## DISCUSSION

Evading the host immune response and reaching immune-privileged organs is essential for *Toxoplasma* to establish a lifelong chronic infection. The complete set of *Toxoplasma* genes involved in these processes is currently unknown. We recently published an *in vivo* loss-of-function screen using a focused sgRNA library enriched in GRA-targeting sgRNAs to determine the *in vivo* fitness contribution of *Toxoplasma* genes in the mouse peritoneum and in distant organs (49). It is likely that a significant fraction of the *Toxoplasma* genes that determined fitness at the site of infection were important for resisting the toxoplasmacidal mechanisms of IFNγ-activated cells or determined parasite growth and survival inside macrophages, the cell type preferentially infected by *Toxoplasma*.

Here, we identified the *Toxoplasma* genes that determine fitness in naïve or IFNγ-activated BMDMs. Many of these parasite genes appear to be involved in nutrient acquisition, likely because IFNγ modulates cholesterol biosynthesis and many other nutrient pathways in macrophages. *Tg*ApiAT3-1, *Tg*ApiAT6-2 or *Tg*ApiAT6-3, which determined fitness in naïve BMDMs, were previously shown to have no fitness defect in HFFs (82) but their phenotype in BMDMs was not investigated. *Tg*ApiAT6-2 was reported to determine resistance to sinefungin (83), which is an S-adenosylmethionine (SAM) analog, likely by mediating the transport of sinefungin into the parasite. SAM is a methyl donor important for many biological processes. Although the *Tg*ApiAT6-2 knockout was reported to have no phenotype *in vitro* or *in vivo,* this was only assessed in the hyper-virulent atypical VAND strain (83). *Tg*ABCG77, which had a lower fitness score in naïve BMDMs *vs*. HFFs, is upregulated by excess cholesterol or LDL and is involved in cholesterol export by *Toxoplasma* (55) suggesting that *Toxoplasma* has a greater need to export cholesterol in BMDMs. It is well known that *Toxoplasma* is a glutton for host-derived lipids and takes up more than it can safely store in lipid droplets. Because excess lipids, especially free cholesterol, can be toxic, our data suggest that in BMDMs, *Toxoplasma* might import more cholesterol and the inability to export excess cholesterol in these cells is likely toxic to the parasite (84). *Toxoplasma* genes involved in DNA repair were more important for fitness in IFNγ-activated *vs*. naïve BMDM. Possibly the generation of oxygen and/or NO radicals generates DNA damage that *Toxoplasma* needs to repair. A previous mutagenesis study (85) identified several *Toxoplasma* mutants that had defects in growth in stimulated murine macrophages. This study identified *Tg*CAP1 (TGGT1_210810) to be important for resistance against NO in macrophages (85). In our study, *Tg*CAP1 had an ∼2-fold larger fitness defect in IFNγ-activated *vs*. naïve BMDMs indicating that *Toxoplasma* genes important for resisting host-generated radicals are important for its fitness. Parasite genes that are important for fitness in IFNγ-stimulated BMDMs were enriched for RNA-binding proteins. Conditions of stress that lead to inhibition of global protein synthesis (e.g., nutrient starvation or oxidative stress) will cause polysomes to disassemble, ribosomes to release their transcripts and the accumulation of mRNA with pre-initiation complexes in stress granules (86). It was previously shown that extracellular *Toxoplasma* have abundant RNA granules, which dissolve when the parasite invades a host cell. The parasites with more abundant “stress granules or RNA granules” have enhanced growth, increased invasion, and decreased apoptosis when extracellularly (87). It is possible that the RNA-binding proteins identified by our screen might play a role in stress granules and adaptation to stress encountered in IFNγ-stimulated BMDM.

Our screen likely missed parasite genes that determine fitness in HFFs but that might have an even larger effect on fitness in naïve or IFNγ-stimulated BMDMs (for example ASP5, (44)) because we started our screen after three to four passages in HFFs. At that point, mutants with a fitness defect in HFFs would have already been depleted and therefore, might not have met the threshold to be included in our analysis (≥ 3 different sgRNAs targeting a specific gene present after the last passage in HFFs). The large number of host cells needed to maintain the complexity of the genome-wide library of parasite mutants made it impractical to directly perform the screen in BMDMs. However, future focused screens could directly start in BMDMs and thereby avoid this limitation. Because IFNγ regulates multiple pathways in murine BMDMs, it is difficult to predict the exact function of each *Toxoplasma* gene that confers fitness in IFNγ-stimulated BMDMs. Comparing the macrophage pathways differentially regulated by IFNγ *vs*. TNFα can start to classify parasite genes that differ in determining fitness in these two conditions. For example, the cholesterol biosynthesis pathway was significantly downregulated in IFNγ-but only marginally downregulated in TNFα-stimulated BMDMs, consistent with previous observations (56), and therefore parasite genes important for adaptation to host cholesterol levels should have fitness defects in IFNγ-but not in TNFα-stimulated BMDMs. However, future secondary screens where only one specific pathway regulated by IFNγ is investigated, for example, the presence of oxygen radicals or the downregulation of the cholesterol pathway, are needed to more definitively classify our hits in different categories.

One of the most significant parasite genes conferring fitness in IFNγ-stimulated BMDMs was GRA45. Deletion of *GRA45* had pleiotropic effects likely because without GRA45, many GRAs aggregated within the parasite’s secretory dense granule organelles. This led to the mislocalization of GRA effectors upon dense granule exocytosis and is also the most likely explanation for its importance in multiple species. GRA16 and GRA24, which are normally secreted beyond the PVM, are no longer secreted in Δ*gra45* parasites. It is unlikely that the fitness defect of Δ*gra45* parasites was due to the defect in GRA effector export beyond the PVM as Δ*myr1* parasites were not significantly more susceptible to IFNγ than wild-type parasites. Although it was previously shown that the secreted GRA effector *Tg*IST is important for resistance against IFNγ, deletion of *Tg*IST had no effect on parasite susceptibility to IFNγ-pre-stimulated cells (36, 37), which is consistent with our results. It is therefore likely that the susceptibility of Δ*gra45* parasites is due to the combined effect of missing multiple secreted effectors and mislocalization of PVM proteins. For example, the PVM-localized GRA23 and GRA17 form the nutrient pore in the PVM and GRA23 was mislocalized in Δ*gra45* parasites. *GRA23* also had a ∼2-fold stronger fitness defect in naïve BMDMs *vs*. HFFs suggesting that *Toxoplasma* might be more reliant on nutrient acquisition *via* PVM nutrient pores in BMDMs. A GRA45 orthologue is exclusively expressed in cat intestinal stages, we speculate that it could be involved in preventing protein aggregation inside wall-forming bodies that are involved in the formation of the oocyst wall.

It was recently shown that phosphorylation of GRAs by WNG1, a closely related family member of WNG2, after exocytosis from the dense granules is important for the correct localization of transmembrane-domain-containing GRAs (68). We have shown that GRA42 and GRA43 also play a role in the correct trafficking of PVM-localized GRAs and that in Δ*gra42* or Δ*gra43* parasites GRA23 and GRA35 are mislocalized (79). Exactly how WNG1, GRA42, GRA43 and GRA45 function in the GRA trafficking pathway is currently unknown. Gram negative bacteria face a similar problem as *Toxoplasma* and also need to transport proteins across several membranes to reach the outer membrane. These bacteria use different periplasmic chaperones and different transport systems to traffick lipoproteins (the Lol system), β-barrel proteins (the Bam system), or Lipopolysaccharides (the Lpt system) (88, 89). It is likely that *Toxoplasma* also has different PV-lumen-localized chaperones and different transport systems to mediate the trafficking of different classes of membrane proteins (e.g., integral *vs*. peripheral *vs*. alpha helical). Possibly genes encoding for some of these were hits in our screen. Future more focussed screens and testing additional knockouts of hits from our screen will allow the identification of *Toxoplasma* genes that mediate the correct trafficking of GRAs to their final destination.

## MATERIALS AND METHODS

### Animals

BMDMs were obtained from 6 to 8 weeks old male/female C57BL/6J mice (Stock No: 000664) and A/J mice (Stock No: 000646) purchased from The Jackson Laboratory. CD-1 female mice were purchased from Charles River Laboratories (Strain Code: 022) and used for *in vivo* infection experiments. 6 to 8 weeks old female/male Brown Norway rats (Strain Code: 091) were purchased from Charles River Laboratories.

### Cell Lines

HFFs were cultured in DMEM, 10% fetal bovine serum (FBS), 2 mM L-glutamine, 100 U/mL penicillin/streptomycin, 10 mg/mL gentamicin. MEFs were cultured in DMEM, 10% fetal bovine serum (FBS), 2mM L-glutamine, 10mM HEPES, 1x non-essential amino acids, 1 mM sodium pyruvate, 100 U/mL penicillin/streptomycin, 10 mg/mL gentamicin. The THP-1 cells were cultured in RPMI-1640, 10% fetal bovine serum (FBS), 2 mM L-glutamine, 100 U/mL penicillin/streptomycin, 10 mg/mL gentamicin. For differentiation, THP-1 cells were stimulated with 100 nM phorbol 12-myristate 13-acetate (PMA) for 3 days and then rested for 1 day by replacing the differentiation medium with complete medium without PMA before performing experiments. Murine BMDMs were isolated from 5 to 8 weeks old female C57BL/6J mice or A/J mice (The Jackson Laboratory) as previously described (90), and obtained by culturing murine bone marrow cells in DMEM, 10% fetal bovine serum (FBS), 2 mM L-glutamine, 10 mM HEPES, 1x non-essential amino acids, 1 mM sodium pyruvate, 100 U/mL penicillin/streptomycin, 10 mg/mL gentamicin and 20% L929 conditioned medium for 5 to 7 days. Rat BMDMs were isolated from 6 to 8 weeks old female Brown Norway rats (Charles River Laboratories) as previously described (49), and obtained by culturing rat bone marrow cells in DMEM, 20% fetal bovine serum (FBS), 2 mM L-glutamine, 10 mM HEPES, 1x non-essential amino acids, 1 mM sodium pyruvate, 100 U/mL penicillin/streptomycin, 10 mg/mL gentamicin and 30% L929 conditioned medium for 5 to 7 days.

### Plasmid construction

The pU6-Universal plasmid (91) was used for generating the plasmid containing sgRNAs targeting the candidate genes. Briefly, the constructs were generated by annealing oligos containing the sequence of sgRNAs (**Table S4**) which were cloned into the BsaI-digested pU6-Universal plasmid. The pUPRT::DHFR-D (Addgene, Cat#58528) plasmid backbone with PCR-amplification to remove the DHFR cassette was used to generate the construct for making the *GRA45* complementation. The promoter region (∼1,500 bp upstream to the start codon) and the coding sequence of *GRA45* were amplified and flanked with the HA epitope sequence before the stop codon. The 3’-UTR region (∼500 bp) was also amplified and assembled with the other two fragments using Gibson Assembly. To construct the plasmid for generating the GRA45 endogenously HA-tagged strain, ∼1,300 bp upstream of the stop codon of GRA45 was amplified by PCR and inserted into pLIC-3xHA-DHFR by ligation-independent cloning (92). All PCR primers are listed in **Table 5S**.

### Generation of parasite strains

*Toxoplasma gondii* parasites were routinely passaged *in vitro* on monolayers of HFFs at 37°C in 5% CO_2_ as previously described (Rosowski et al., 2011). Individual knockout of candidate genes was performed using CRISPR-Cas9 (91, 93). To generate the knockout strains for candidate fitness-conferring genes in IFNγ-activated BMDMs, plasmids containing sgRNAs were co-transfected with NotI (New England Biolabs)-linearized pTKO, which contains the *HXGPRT* selection cassette and GFP (94), into RH-Luc+/Δ*hxgprt* parasites at a ratio 5:1 of sgRNA to linearized pTKO plasmid. 24 h post-transfection, populations were selected with mycophenolic acid (50 μg/mL) and xanthine (50 μg/mL) and cloned by limiting dilution (**Figure S2A**). Individual knockout clones were confirmed by PCR (**Figure S2D to S2F**). In addition to generating knockouts in the RH-Luc+/Δ*hxgprt* strain, the RH-Cas9/Δ*hxgprt* strain (95) was used to generate Δ*gra22* and Δ*rom1* parasites (**Figure S2G and S2H**) using the same strategy. To generate the *GRA45* complemented strain (**Figure S2B**), the RH-Luc+/Δ*gra45* parasites were co-transfected with plasmids containing sgRNAs specifically targeting the *UPRT* locus and SalI (New England Biolabs)-linearized pUPRT::GRA45HA plasmid at a ratio 1:5 of sgRNAs to linearized plasmid. After the first 2 complete lysis cycles, populations were selected with 10 μM of 5-fluoro-2-deoxyuridine (FUDR) for another 2 complete lysis cycles. Individual clones were isolated by limiting dilution and the presence of *GRA45HA* was determined by immunofluorescence assay (IFA) and by PCR to confirm the integration into the *UPRT* locus (**Figure S2D**). The *GRA45* knockout in RHΔ*ku80*::*MYR1*-*3xHA* strain (kindly provided by Dr. John Boothroyd) was generated by co-transfecting plasmids containing sgRNAs targeting the *GRA45* locus along with an amplicon harboring *GRA45* homology regions (60 bp) surrounding a pyrimethamine-resistant (DHFR*) cassette (**Figure S2C**). Individual knockout clones were grown in medium supplemented with 3 μM pyrimethamine followed by limiting dilution and subsequent screening by PCR for correct integration of *DHFR\** into the *GRA4*5 locus (**Figure S2I**). GRA45 endogenously HA-tagged parasites were made in the RH*Δku80Δhxgprt* strain (92) by transfection with the plasmid pLIC-GRA45-3xHA-DHFR. Transfected parasite populations were selected with 3μM pyrimethamine and cloned by limiting dilution. The presence of the *GRA45*-*3xHA* was determined by immunofluorescence assays.

### *Toxoplasma gondii* CRISPR-Cas9 mediated genome-wide loss-of-function screens

At least 5×10^8^ parasites were transfected with a mixture of pU6-DHFR plasmids (100 μg for each 1×10^8^ parasites) containing 10 different sgRNAs against every *Toxoplasma* gene. A monolayer of human foreskin fibroblasts (HFFs) was subsequently infected with the parasites (MOI=0.5) and grown for 24 h in DMEM containing 1% FBS, 2 mM L-Glutamine, 1% penicillin/streptomycin, and 40 µM Chloramphenicol (CAT). Subsequently, the medium was removed and replaced with DMEM containing 10% FBS, 2 mM L-Glutamine, 10 mM HEPES, 1x Non-Essential Amino Acids, 1 mM Sodium Pyruvate, 100 U/mL Pen/Strep, 10 µg/mL gentamicin, 40 µM CAT Chloramphenicol, 1 µM Pyrimethamine, and 10 µg/mL DNase I for 3 or 4 passages. To perform the screen in murine macrophages, BMDMs isolated from C57BL/6J mice were stimulated with: 100 ng/mL IFNγ for 24 h for screen 1 (S1), 1 ng/mL IFNγ for 24 h for screen 2 (S2), or 100 ng/mL IFNγ for 4 h or 100 ng/mL TNFα for 4 h for screen 3 (S3). Both naïve BMDMs and stimulated BMDMs were infected with the mutant pool derived from the 3^rd^ or 4^th^ passage in HFFs at an MOI of 0.5. After 48 h infection (the time the mutant parasites nearly egressed from the host cells), the parasites were harvested, counted and used to infect macrophages that had undergone the same stimulation until the 3^rd^ passage (**Figure 1B**). For each passage a pellet of 1×10^7^ parasites were collected and used for genomic DNA extraction and PCR amplification of the sgRNA with a barcoding primer. The sample was sent for Illumina sequencing at the University of California Davis Genomic Center on a NEXT Seq (Illumina) with single-end reads using primers (P150 and P151) listed in **Table S4**.

### RNA-seq

3×10^6^ BMDMs were seeded overnight in 6-well plates prior to stimulations. For the stimulated samples, IFNγ (100 ng/mL) and/or TNFα (100 ng/mL) were added to each well for 4 or 24 h before harvesting the cells for total RNA extractions. The RNA-seq libraries were prepared, sequenced, and processed as previously described (96). Briefly, mRNA was purified by polyA-tail enrichment (Dynabeads mRNA Purification Kit, Thermo Fisher), fragmented into 200–400 base-pairs, and reverse transcribed into cDNA before adding Illumina sequencing adapters to each end. Libraries were barcoded, multiplexed into 4 samples per sequencing lane in the Illumina HiSeq 2000, and sequenced from both ends resulting in 50 bp reads after discarding the barcodes. The RNA-sequencing reads were mapped to the mouse genome (mm10) using Bowtie (2.0.2) (97) and Tophat (v2.0.4) (98) and transcript abundance estimated in cufflinks (99).

### Bioinformatic analysis of the loss-of-function screens and RNA-seq

sgRNA selection and screen analysis were performed using custom software as described in (Sidik et al., 2016, 2018) that will be provided upon request. Statistical analyses were performed in R (www.R-project.org) and Excel (Microsoft Office). Illumina sequencing reads were matched against the sequences of the sgRNA library. The number of exact matches was counted and considered as raw read numbers. The abundance of each sgRNA was normalized to the total number of reads. To do the log2 transfer, sgRNAs that had zero reads were assigned a pseudo-count corresponding to 0.9, which is 90% of the lowest sgRNA read in the sample. Only sgRNA whose abundance was above the 5th percentile in the input (library) were further considered for the analysis. The ‘‘fitness’’ for each gene was calculated as the average log2 fold change for the top five scoring sgRNAs, which minimized the effect of stochastic losses. The Pearson correlation between each sample from different screens was calculated. To identify the genes that underwent negative selection, the raw reads of each sgRNA between two samples was compared and the negative selection *P*-value of each gene was calculated using the MAGeCK algorithm (50). Genes were considered fitness-conferring if they met a significance threshold of negative selection *P* < 0.05 in two out of three screens and/or ≥ 2-fold lower fitness (the fitness between groups is ≤ -1.0) in two out of three screens.

Gene set enrichment analysis (51) was used to determine if *Toxoplasma* genes that determine fitness in specific conditions were enriched in the functional annotation. The genes with fitness-conferring in specific conditions were analyzed by using an in-house database that contained information on gene ontology (GO), protein family domains (Interpro), KEGG enzyme EC numbers, localization to specific organelles, amongst others. Pathways that were enriched in at least two screens are indicated in **Table S3**. For enrichment analysis of BMDM pathways stimulated with IFNγ *vs*. unstimulated or IFNγ *vs*. TNFα, the GSEA program and MsigDB database was used (51, 100, 101). Psi-blast was used to find orthologues of proteins under investigation in other species. Alignments of GRA45 with its orthologs were made using PRALINE (102), and the results (**Figure S4A**) were used as input for HHpred (103) to predict similarity to secondary structures of other proteins. In addition, the SWISS-MODEL server was used for the homology modeling of protein structures (104).

### Plaque assay in MEFs

Freshly confluent 24 well plates of MEFs were infected with 100 parasites for each well and the plates were incubated at 37°C without disturbing for 5 days. The area of each individual plaque was captured and analyzed using a Nikon TE2000 inverted microscope equipped with Hamamatsu ORCA-ER digital camera, and NIS Elements Imaging Software respectively. For each independent experiment, at least 40 plaques from technical duplicate wells were imaged.

### Luciferase assay

Luciferase assays were performed to determine the fitness of knockout strains in IFNγ-activated macrophages. In murine and rat macrophages, BMDMs in 96-well plates (1×10^5^ cells/well) were stimulated with IFNγ (1 ng/mL or 5 ng/mL) for 24 h followed by infection with wild-type luciferase-expressing parasites or knockout parasites at 2 different MOIs (MOI = 0.5 and 1). For human macrophages, differentiated THP-1 cells in 96-well plates (1×10^5^ cells/well) were stimulated with human IFNγ (50 U/mL or 500 U/mL) for 24 h followed by parasite infection. A plaque assay was set up in HFFs at the same time to determine the parasite viability of each parasite strain. After 24 h infection, the cells were lysed and luciferase activity was measured as previously described (105). Raw luciferase reads (RLU) of unstimulated infected cells was considered as 100 percent and relative parasite growth in IFNγ-activated macrophages was calculated. To make comparisons between wild-type and knockout parasites, “real” MOI was matched to 0.25 from the plaque assay results.

### Growth competition

The BMDMs were left unstimulated or stimulated with IFNγ (1 ng/mL) for 24 h followed by infection with a 1:1 mixed ratio of GFP-expressing wild-type parasites and GFP-negative knockout parasites (Δ*gra22* or Δ*rom1*). The mixed parasites were allowed to grow in the BMDMs for three lytic cycles. Plaque assays of the mixed parasites were performed before putting into the BMDMs and after the 3^rd^ passage in the BMDMs, and the number of GFP positive *vs*. GFP negative plaques were counted to determine the ratio of wild type and knockout parasites in naïve BMDMs and IFNγ-activated BMDMs.

### Immunofluorescent assay

To check the localization of GRA45 in extracellular parasites, RHΔ*ku80*Δ*hxgprt::GRA45-3xHA* tachyzoites released from syringe-lysed HFFs were loaded onto coverslips and fixed with 4% Paraformaldehyde (PFA) for 20 minutes followed by permeabilization/blocking with PBS with 3% (w/v) BSA, 5% (v/v) goat serum and 0.1% Triton X-100. Co-localization of GRA45 was detected by anti-HA antibodies along with antibodies against ROP2/3/4, GRA1, GRA2, GRA5, GRA7 or SAG1. To determine the localization of intracellular GRA45, HFFs grown on glass coverslips were infected with RHΔ*ku80*Δ*hxgprt::GRA45-3xHA* parasites for 24 h, fixed with 4% PFA for 20 minutes, permeabilized/blocked with PBS with 3% (w/v) BSA, 5% (v/v) goat serum and 0.1% Triton X-100, followed by incubated with antibodies against the HA epitope tag together with antibodies against ROP2/3/4, GRA1, GRA2, GRA5, GRA7 or SAG1 at 4°C overnight. To check the PVM localization of GRA23 or export of GRA16 and GRA24, HFFs grown on glass coverslips were infected with wild-type, Δ*myr1,* Δ*gra45* or Δ*gra45+GRA45HA* strains in the RHΔ*hxgprt::cLuc* background transiently expressing GRA23-HA-FLAG, GRA16-HA-FLAG or GRA24-HA-FLAG for 20-24 h followed by fixation, permeabilization and blocking as described above. The coverslips were incubated with anti-SAG1 antibodies and anti-HA antibodies at 4°C overnight. Alexa Fluor 488/594 secondary antibodies and DAPI were used as previously described (79). The coverslips were mounted with Vecta-Shield mounting oil and the microscopy was performed with NIS-Elements software (Nikon) and a digital camera (CoolSNAP EZ; Roper Scientific) connected to an inverted fluorescence microscope (Eclipse Ti-S; Nikon) and either phase contrast or DIC imaging.

### Cell fractionation

Purified extracellular parasites were washed with PBS and resuspended in cold PBS containing 1x protease and phosphatase inhibitors (Thermo Fisher). Parasites were lysed by six to eight freeze/thaw (F/T) cycles using a dry ice-ethanol bath (-70°C) and a 37°C bath. 1/3 of the F/T lysate was kept in PBS, 1/3 in 1% NP-40, and 1/3 in 1% Triton X-100 for 30 minutes at 4°C under rotation. The PBS, NP-40, and Triton X-100 treated lysates were separated by high-speed spin 30 minutes at 20,000 *× g*. The PBS supernatant was precipitated with TCA (Trichloroacetic Acid), NP-40, and Triton X-100 supernatants were acetone precipitated. For intracellular parasites, tachyzoites were allowed to infect HFF monolayers for 24 h. The infected monolayers were washed three times with PBS, mechanically lysed in PBS containing 1x protease and phosphatase inhibitors (Thermo Fisher) using 27-gauge needles, and parasites were pelleted by low-speed spin for 10 minutes at 2500 × *g* (LSP). The low-speed supernatant (LSS) containing PV membranes and soluble material was further fractionated at 100,000 *x g* at 4°C for one hour to yield the high-speed pellet (HSP) and high-speed supernatant (HSS). Half of the HSP was treated with 1% Triton X-100 for 30 minutes at 4°C under rotation and fractionated again at 20,000 *x g* at 4°C for one hour to obtain the Triton X-100 treated high-speed pellet and high-speed supernatant.

### Immunoblotting

Samples from the extracellular and intracellular fractionations were boiled for 10 minutes in sample buffer, separated by SDS-PAGE, and transferred to polyvinylidene difluoride (PVDF) membranes. Membranes were blocked in 5% milk in TBS supplemented with 1% Tween-20 (TBS-T) for 30 minutes at room temperature and then incubated overnight at 4°C with primary antibody in blocking buffer. GRA1, GRA2, GRA5 blots were subsequently incubated with an anti-mouse secondary horseradish peroxidase (HRP)-conjugated antibody (Roche). The GRA7 blot was incubated with a secondary antibody (HRP)-conjugated anti-rabbit antibody (Roche). The HRP was detected using an enhanced chemiluminescence (ECL) kit (Pierce). Between each blot, the membranes were stripped by incubation in stripping buffer (Thermo Fisher) for 20 minutes and subsequently washed three times for five minutes with TBS-T.

### *In vivo* infection

*Toxoplasma* tachyzoites were harvested from cell culture and released by passage through a 27-gauge needle, followed by a 30-gauge needle. C57BL/6J mice were intraperitoneally (i.p.) infected with 100 tachyzoites of each strain and parasite viability of the inocula was determined in a plaque assay after infection. The mice were monitored for 30 days p.i., and the number of dead mice per group was observed every individual day.

### Statistical Tests

All statistical analyses were performed using Prism (GraphPad) version 8.0. The luciferase assays are presented as paired scatterplots for independent experiment whereas all the other data are presented as average ± standard deviation (SD), and the exact n values are mentioned in the figure legends. For all the calculations p < 0.05 are considered as significant and represented with an Asterix. To compare parasite growth of the knockouts *vs*. wild-type parasites in IFNγ-activated cells, paired *t* test was used. For more than three groups with one variable in the other data, One-way ANOVA with Tukey’s multiple comparisons test was used. For one variable test with two groups, the two-way ANOVA with Tukey’s multiple comparisons test was used. Survival experiments were analyzed using the log rank (Mantel-Cox) test.

## ACKNOWLEDGEMENTS

This study was supported by the National Institutes of Health (R01-AI080621) awarded to J.P.J.S. NIH’s Director’s Early Independence Award (1DP5OD017892), and a Mathers Foundation grant to S.L. We acknowledge all members of the EupathDB.org team for generating this invaluable resource without which this work would not have been possible.

## AUTHOR CONTRIBUTIONS

J.P.J.S., Y.W., and LO.S. designed experiments and wrote the manuscript with input from all authors. Y.W. and LO.S. performed and interpreted most of the experimental work. T.C.PS., S.K. helped with the CRISPR screen. M.H. performed the RNAseq experiments. A.M.F. helped with the generation of knockout parasites and growth competition assays. B.M.M. generated the RH-Cas9Δ*hxgprt* strain. S.L. contributed reagents and analysis tools and provided input on the design of the experiments.

## SUPPLEMENTAL INFORMATION

**Supplemental Table S1**

<u>Sheet 1</u> “BMDM_AJ_B6_IFNGvsTNFvsIFNTNF_FP”: RNAseq data of A/J and C57BL/6J BMDMs stimulated with IFNγ and/or TNFα for 4 or 24 h or left unstimulated. RPKM values are indicated. Genes involved in the cholesterol synthesis pathway are indicated as are pathways mediating cholesterol biosynthesis.

<u>Sheet 2</u> “GSEA_IFN4-24hvsTNF4-24h”: Gene Set Enrichment Analysis (GSEA) was performed on the BMDMs stimulated with IFNγ *vs*. TNFα. For this analysis the A/J, C57BL/6J at 4 and 24 h were treated as four replicates to identify pathways regulated regardless of mouse strain or duration of stimulation. All data in sheet 1 was used and genes were ranked based on the differential expression between the two conditions identified using the Signal2Noise option in the stand-alone version of GSEA. KEGG and Hallmark gene sets from the MSigDB database were used to identify positively or negatively enriched gene sets. A *P*-value cut-off of 0.05 was used but FDRs are also displayed. 1,000 gene set permutations were used to determine *P*-values and FDRs.

<u>Sheet 3</u> “GSEA IFN4-24 vs NS4-24”: As in “Sheet 2” but comparing BMDMs stimulated with IFNγ *vs*. non-stimulated (NS) BMDMs.

<u>Sheet 4</u> “GSEA TNF4-24vs NS4-24”: As in “Sheet 2” but comparing BMDMs stimulated with TNFα *vs*. non-stimulated (NS) BMDMs.

**Supplemental Table S2**

This table contains the raw sgRNA reads for each experiment, the fitness scores, and a list with all *Toxoplasma* genes and all associated screen information.

**Supplemental Table S3**

Gene Set Enrichment Analysis (GSEA) using the pre-ranked option was performed on *Toxoplasma* genes, with at least 3 sgRNAs in either condition, ranked according to their Naive BMDM *vs*. HFF-P6/8 *P*-values or according to their IFNγ BMDM *vs.* Naive BMDM *P*-values (see Supplementary Table S2). Only gene sets containing at least five genes were analyzed and only those gene sets enriched in at least 2 out of the 3 screens with in at least one screen a *P*-value <0.05 are listed. For enrichments mentioned in the main text but not listed in this table the “results analysis” option in ToxoDB.org on the top hits was used.

**Supplemental Table S4**

Primers and oligos used in this study.

---

**Figure S1.**
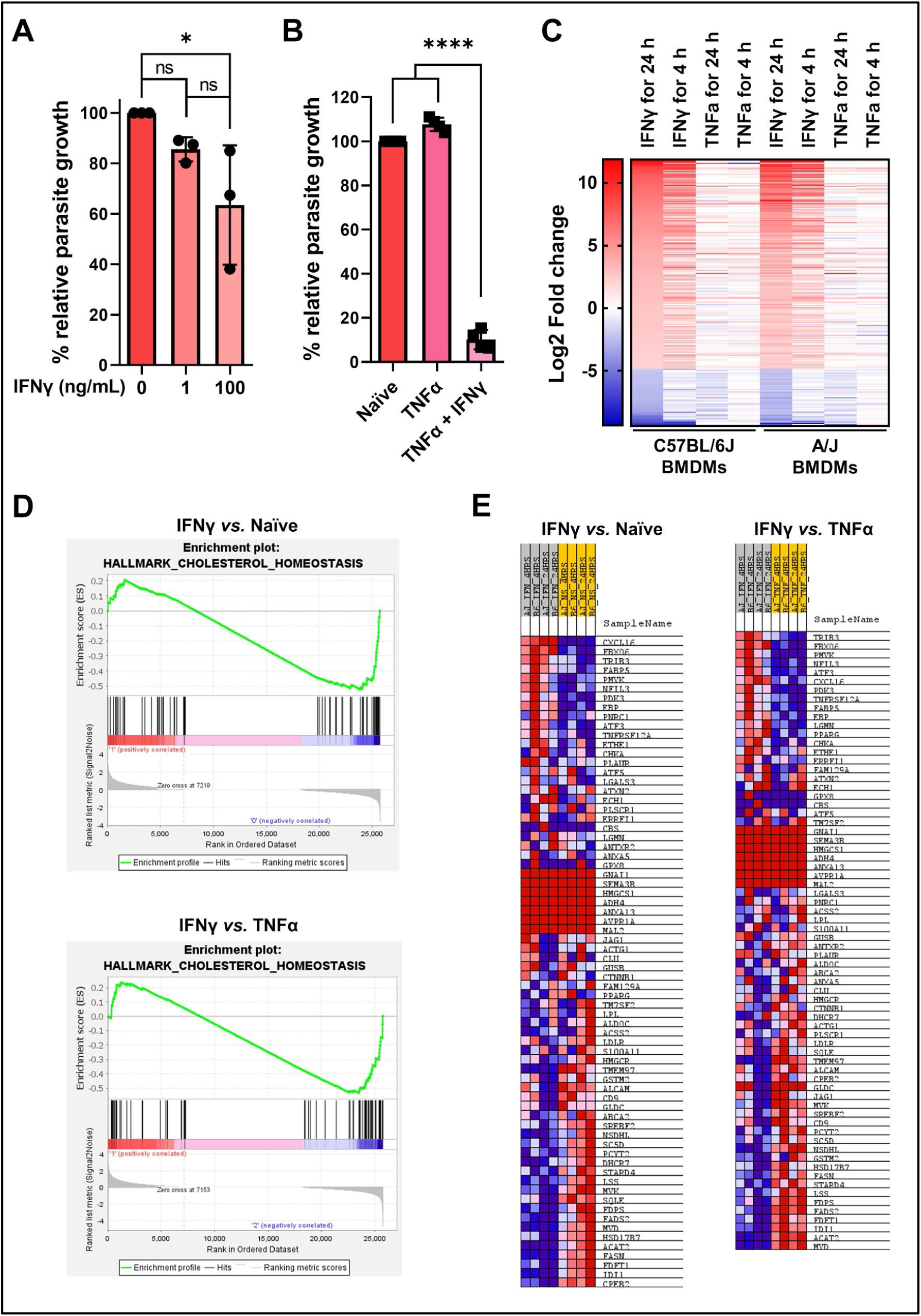
IFNγ restricts parasite growth in murine BMDMs, related to Figure 1. (A) Murine BMDMs pre-stimulated with IFNγ (1 or 100 ng/mL) or left unstimulated for 24 h were infected with luciferase-expressing type 1 (RH) parasites for 24 h and parasite growth was measured by luciferase assay. Parasite growth in IFNγ-activated BMDMs is expressed relative to growth in naïve BMDMs. Data are displayed as average ± SD for 3 independent experiments. Asterisk (*) indicates a significant difference analyzed with one-way ANOVA with Tukey’s multiple comparisons test: *p* = 0.04 for IFNγ (100 ng/mL) stimulated BMDMs *vs.* naïve BMDMs. (B) Murine BMDMs pre-stimulated with IFNγ (100 ng/mL) and/or TNFα (10 ng/mL) for 24 h or left unstimulated were infected with luciferase-expressing RH parasites for another 24 h and parasite growth was measured by luciferase assay. Parasite growth in stimulated BMDMs is expressed relative to growth in naïve BMDMs. Data are displayed as average ± SD for 4 independent experiments. Asterisk (*) indicates a significant difference analyzed with one-way ANOVA with Tukey’s multiple comparisons test: *p* < 0.0001 for TNFα + IFNγ stimulated BMDMs *vs.* both naïve BMDMs and TNFα stimulated BMDMs. (C) IFNγ-regulated genes in C57BL/6J and A/J murine BMDMs pre-stimulated with 100 ng/mL IFNγ were defined by a ≥ 4-fold change (IFNγ-activated BMDMs *vs.* naïve BMDMs) in gene expression values (FKPM). Data are displayed as a heat map of log_2_-fold change of the 251 genes with either ≥ 4-fold upregulated (197 genes) or ≥ 4-fold downregulated (54 genes) in IFNγ-activated BMDMs (*n* = 2). The complete set of genes is listed in **Table S1**. (D) Differentially regulated pathways between unstimulated *vs*. IFNγ-*vs*. TNFα-stimulated BMDMs were identified using GSEA analysis and the MSigDB database. The 4 and 24 h time points from A/J and C57BL6/J murine BMDMs were treated as replicates as the goal was to identify pathways that are similarly regulated between these samples. As an example, the downregulation of the cholesterol homeostasis pathway in IFNγ-activated vs. Naive BMDMs and IFNγ-activated vs. TNFα-activated BMDMs are shown. (E) An example of specific genes involved in the cholesterol homeostasis pathway (from D) regulated in IFNγ-stimulated murine BMDMs are presented as heat plots. Exact RPKM values of genes involved in cholesterol metabolism is presented in **Table S1**.

**Figure S2.**
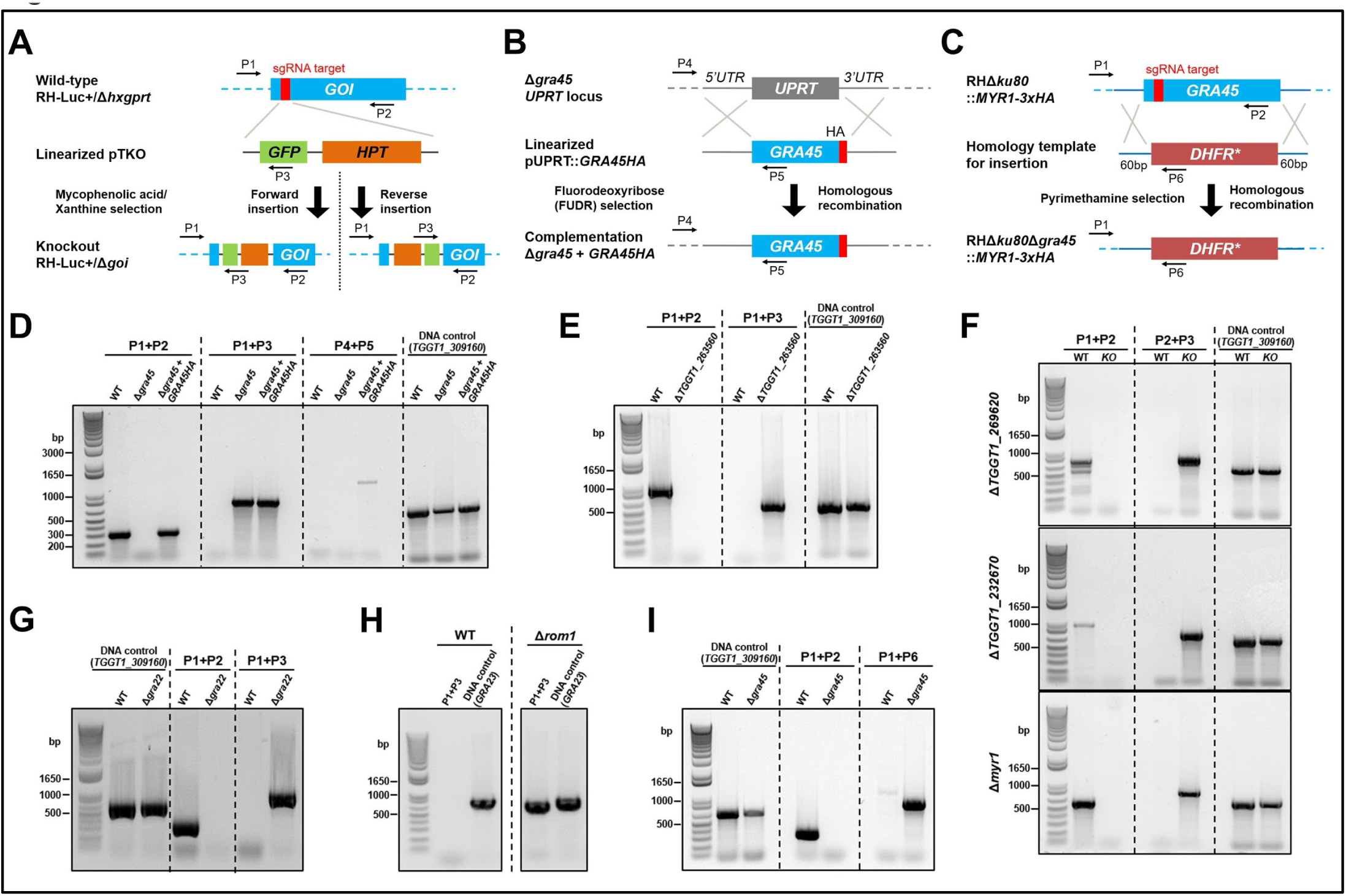
Generation and confirmation of knockout and complemented parasites, related to Figure 2. (A) Schematic diagram depicting the genomic loci of the genes of interest (GOI) (top) and the CRISPR/Cas9-targeting site (red box). Linearized pTKO plasmid containing GFP-coding sequence and HXGPRT selection cassette (middle) was used as a repair template to disrupt the GOI (bottom) after mycophenolic acid and xanthine selection. P1 and P2 refer to the primers used for confirming locus disruption. P1 + P3 or P2 + P3 are used to check insertion of the repair template into the GOI locus. (B) GRA45 complementation was performed by homologous recombination of *GRA45HA*-expressing cassette (middle) into *UPRT* locus (top). The coding sequence of *UPRT* was replaced with *GRA45HA* (bottom) after FUDR selection. (C) Schematic strategy used to delete the entire coding region of GRA45 (top) by inserting DHFR* in RHΔ*ku80*::*MYR1*-*3xHA* strain. Transfection of the sgRNAs targeting *GRA45* locus (red box) together with the DHFR*-expressing amplicon flanking with 60bp of 5-’UTR and 3’-UTR of GRA45 (middle) after pyrimethamine selection was used to generate *GRA45* deletions. (D) *GRA45* knockout in RH-Luc+/Δ*hxgprt* parasites and complementation of the gene in the *UPRT* locus. (E) *TGGT1_263560* knockout in RH-Luc+/Δ*hxgprt* parasites. (F) *TGGT1_269620* (top), *TGGT1_232670* (middle) and *MYR1* (bottom) knockout in RH-Luc+/Δ*hxgprt* parasites. (F and G) Individual knockout of *GRA22* (F) or *ROM1* (G) in RH-Cas9Δ*hxgprt* background. (I) *GRA45* knockout in RHΔ*ku80*::*MYR1*-*3xHA* parasites.

**Figure S3.**
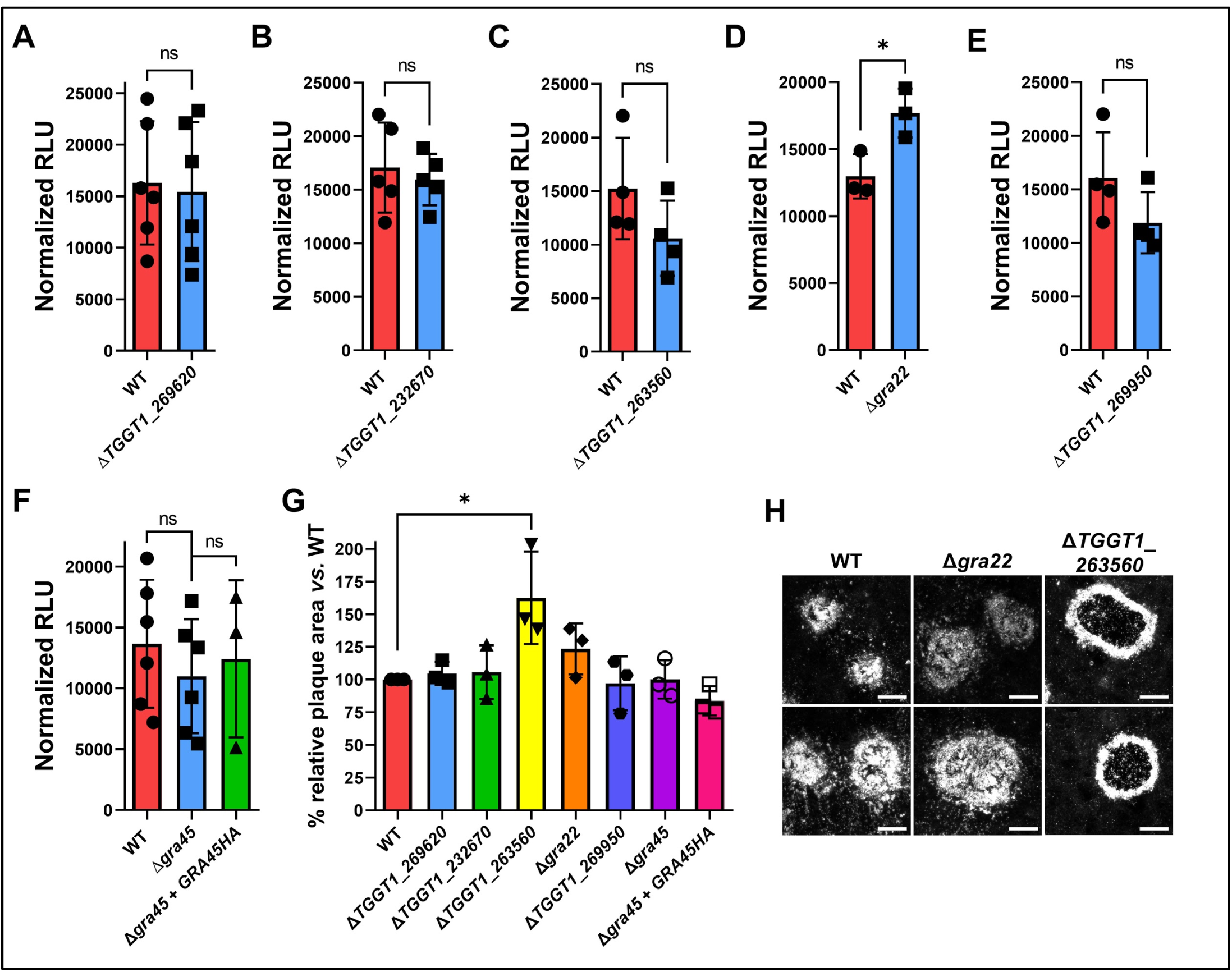
Parasite growth in naïve murine BMDMs and MEFs of individual knockouts of genes identified from loss-of-function screen in IFNγ-stimulated BMDM, related to Figure 2. (A to F) Raw luciferase reads (RLU) of WT, Δ*TGGT1_269620* (A, *n* = 6), Δ*TGGT1_232670* (B, *n* = 5), Δ*TGGT1_263560* (C, *n* = 4), Δ*gra22* (D, *n* = 3), Δ*TGGT1_269950* (E, *n* = 4) or Δ*gra45 (n* = 6) and Δ*gra45*+*GRA45HA (*F, *n* = 6 for Δ*gra45*, *n* = 3 for Δ*gra45*+*GRA45HA*) parasites in naïve murine BMDMs were normalized to 100% viability based on their viability measured by plaque assays. Data are displayed as averages ± SD for at least 3 independent experiments. Asterisk (*) indicates a significant difference between WT and knockout analyzed with two-tailed paired *t* test: *p* = 0.01 for (D). (G) Confluent MEFs were infected with indicated parasites for 5 days. An area of at least 40 plaques per experiment was measured. Data are displayed as averages ± SD for 3 independent experiments with plaque area relative to the WT plaque area (*n* = 3). Asterisk (*) indicates a significant difference analyzed with one-way ANOVA with Tukey’s multiple comparisons test: *p* = 0.02 for WT *vs.* Δ*TGGT1_263560*. (H) Representative images of plaque size and morphology of WT, Δ*gra22*, and Δ*TGGT1_263560* parasites in MEFs (scale bar = 200 μm).

**Figure S4.**
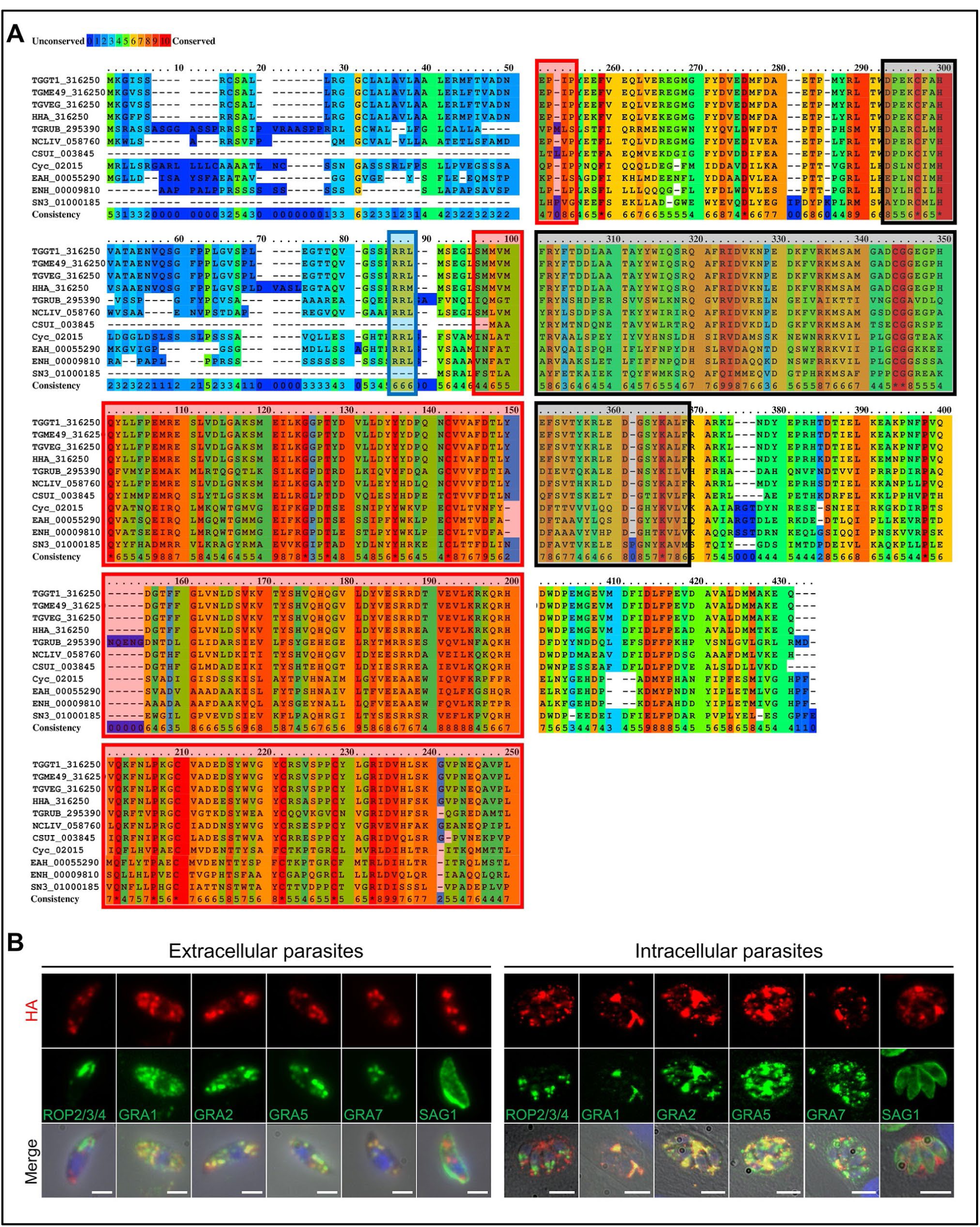
The subcellular localization and predicted structure of GRA45. Related to Figure 4. (A) Alignment of GRA45 from *Toxoplasma gondii* type 1 (TGGT1_316250), 2 (TGME49_316250) and 3 (TGVEG_316250); *Toxoplasma gondii* TGRUB_295390; *Hammondia hammondi* HHA_316250; *Neospora caninum* NCLIV_058760; *Cystoisospora suis* CSUI_003845; *Sarcocystis neurona* SN3_01000185; *Eimeria acervulina* EAH_00055290; *Eimeria necatrix* ENH_00009810; *Cyclospora cayetanensis* cyc_02015. Blue box indicates TEXEL motif. Red boxes and black boxes indicate predicted small heat shock protein (HSP) domain and predicted transthyretin-like fold, respectively. (B) Extracellular parasites (left panel) or intracellular parasites (right panel) expressing endogenously HA-tagged GRA45 were fixed, permeabilized, and subjected to immunofluorescent assays with the indicated antibodies. (scale bar = 2 μm for extracellular and 5 μm for intracellular parasites).

